# Binding dynamics shape germinal center broadly neutralizing responses to HIV priming

**DOI:** 10.64898/2026.05.12.724749

**Authors:** Jinsong Zhang, Jim Sindayen, Miyo Ota, Kara Anasti, Zbignew Mikulski, Angel Gandarilla, Takayuki Ota, Daniel Ramirez, Amanda Zhang, S. Munir Alam, Marilyn Diaz, Laurent Verkoczy

## Abstract

Inducing broadly neutralizing antibodies (bnAbs) is central to HIV vaccine efforts, but bnAb precursors are rare, often inactive, and require extensive somatic hypermutation (SHM) to recognize diverse, glycan-shielded epitopes. Germline-targeting immunogens (GTs) aim to jump-start this process, but determinants of success remain unclear. Here we establish a bnAb precursor-trackable model that reveals a surprising driver: binding dynamics. Across a series of GTs, multivalent designs that engage B-cells transiently - not tightly - consistently outperformed others, boosting germinal center fitness and unlocking rare or more efficient SHM pathways for breadth. These effects were independent of affinity or precursor frequency. Single-cell transcriptomics uncovered gene programs that predict successful priming. Crucially, this scalable system provides general predictive power for immunogen performance, marking a major advance for not only HIV vaccine development, but also potentially establishing a broadly applicable framework for streamlining pre-clinical pipelines, including immunogenicity and safety evaluation.

## INTRODUCTION

A vaccine capable of preventing HIV infection remains a global priority^1^. A central obstacle is the induction of broadly neutralizing antibodies (bnAbs) against relatively conserved regions of the highly glycosylated and diverse HIV-1 envelope glycoprotein (Env)^2–5^. Rare bnAb precursor clonotypes isolated after prolonged infection have revealed their immunogenetics, mode of Env recognition, and evolutionary pathways. Collectively, this indicates that bnAb induction lies at the intersection of structural constraints and immunoregulation: recognizing Env’s well-concealed epitopes requires bnAb clonotypes with unusual features—including long or flexible HCDR3s^6^, restricted germline usage^7^, and/or auto-/polyspecificity^8–11^. These properties render bnAb precursors rare in the repertoire and subject to peripheral tolerance mechanisms like clonal anergy, diminishing B-cell Receptor (BCR) signaling capacity^12–14^.

These atypical precursor features, combined with the unusual immunobiology of HIV-1 as a rapidly integrating retrovirus^7,15^, sharply distinguish HIV vaccination from conventional immunization^16^. Whereas most vaccines mobilize abundant, fully competent clonotypes requiring only modest somatic hypermutation (SHM) within short-lived germinal centers (GCs), HIV vaccination must recruit rare, often tolerance-constrained precursors amid competition from immunodominant clones. Three challenges follow. First, bnAb precursors must be selectively recruited and rapidly activated in GCs to achieve protection before early viral entry. Second, sustained GC residence with iterative dark zone–light zone cycling (or GC recycling) is required to sufficiently expand mutational landscapes to attain improbable point mutations or insertions/deletions^17–19^ and support efficient recall. Such rare SHM events enhance breadth in clonotypes with partial germline Env recognition (e.g., V2 apex–directed bnAbs) and are essential for others that initially lack Env specificity, such as CD4 binding site–targeting VRC01/CH31-class precursors expressing the hypermutable V_H_1-2 gene segment^20–22^. Third, extended GC persistence may be necessary to extinguish self-reactivity while refining broad Env recognition through clonal redemption^23–27^, a purifying selection process that, in HIV, may require multiple rounds^28–31^. Thus, an effective HIV vaccine must couple rapid SHM initiation with sustained, diversified GC evolution capable of generating breadth within a practical immunization regimen—demands far exceeding those for conventional pathogens.

To overcome the earliest bottleneck in bnAb induction, germline-targeting (GT) immunogens were developed to selectively engage inferred germline or unmutated common ancestor (UCA) precursor BCRs at priming^19,32–41^. The most extensively optimized platform, the eODGT scaffold, culminated in the eODGT8 60-mer, reverse engineered to bind VRC01/CH31-class precursors with ultra-low (low nM) affinity^33–35^. Several GT immunogens, including eODGT8, have advanced to early clinical trials^42–44^, and reliably recruit and expand targeted precursors in humans, yet induce limited SHM. Moreover, incorporation into current protocols has yielded only sporadic, transient, or narrow serum cross-neutralizing responses, even after multiple boosting steps^19,45–47^—not markedly distinct from non-GT approaches^48–50^. A prevailing posit is that high equilibrium affinity (K_D_) is required to offset low precursor frequency and ensure competitive GC entry^51–54^. However, some studies show an impact on clonal expansion rather than GC fitness^55^, while classical B cell studies (albeit, performed with abundant clonotypes) suggest moderate affinity is optimal, implying an affinity “ceiling” ^56–58^. Because K_D_ reflects the ratio of association and dissociation rates, binding kinetics may independently shape receptor clustering, signaling thresholds, and ultimately, fate decisions^59,60^. In this context, we recently showed that proximal activation of VRC01-class precursors depends on achieving a minimal association rate rather than K_D_, suggesting that kinetic thresholds govern early BCR signaling^61^. Whether such biophysical parameters control the rate-limiting step of GC recruitment and maturation *in vivo*, particularly under host controls and with clonal competition, remains unresolved.

BnAb knock-in (KI) models have become essential for addressing these questions^62,63^. Conventional models, including wild-type mice and macaques, often lack homologues of human bnAb precursors, including the V_H_1-2–restricted VRC01/CH31 class, necessitating KI systems for preclinical evaluation^64^. Furthermore, KI→WT chimeras, in which genetically marked KI precursors are introduced into polyclonal repertoires at defined frequencies, permit controlled tracking of their activation, clonal expansion, and GC recruitment^53–55,65^. Such transfer studies have shown higher order eODGT variants enhance V_H_1-2 restricted precursor fitness in GCs, particularly at physiologic frequencies^65^. Beyond practical testing, KI models also provide the resolution to dissect lineage-specific pre- and post-immunization bottlenecks and refine vaccine strategies. For example, comparisons across KI models expressing distinct V_H_1-2-restricted V(D)J rearrangements have revealed that even closely related clonotypes exhibit variable peripheral anergy^19,35,66–68^ and that for more profoundly unresponsive ones, GT multimerization is crucial for their clonal expansion^66^, akin to the prototype, 2F5 bnAb anergy model^28,69^.

For physiologic and technical reasons, V_H_1-2–restricted CH31 UCA dKI animals^19^ offer a particularly tractable model to link BCR sensing with *in vivo* GC programming. Its main vaccine target (follicular B-cells), express the authentic human CH31–CH34 lineage UCA clonotype (V_H_1-2D_H_3-16J_H_4/ Vk1-33 J_k_2) BCRs which, at baseline, functionally resemble mildly polyreactive ones having a competitive GC fitness advantage^23,24,70–73^. Additionally, its knocked-in V_H_1-2 minigene is intrinsically hypermutable, and the clonotype exhibits a (so-far) unique propensity to acquire breadth-associated heavy chain insertions upon immunization, thereby adding practicality and stringency to evaluate SHM quality. In this study, we generate CH31 UCA^hom/hom^ ^dKI^→WT chimeras and prime them with closely related eODGT variants (differing in affinity, kinetics, and valency) to systematically define how discrete biophysical parameters shape GC recruitment, cycling, and persistence *in vivo* in relation to proximal BCR signaling *ex vivo*. We also establish a complementary CH31 UCA^hom/hom^ ^dKI^ culture platform to interrogate GC transcriptional programs and, in so doing, link proximal signaling and *in vivo* priming thresholds, ultimately defining a potential high-throughput, predictive *ex vivo* modality for testing antigen-specific GC priming quality.

## RESULTS

### Effect of GT immunogen affinity (K_d_), binding kinetics (k_a_/k_d_) and valency on proximal bnAb precursor signaling *ex vivo*

We previously showed, using Env proteins with varying affinities and kinetic binding rates for ‘CD4 mimic’ bnAb-class BCRs, that minimal association rate (k_a_), rather than overall affinity (K_D_), governs proximal B-cell activation ^61^. However, these studies relied on Ramos B-cell lines, which differ substantially from nascent mature follicular (MF) B-cells—the principal vaccine targets undergoing GC recruitment in mice and humans. These differences include: (i) signaling thresholds of BCRs expressed as IgGs rather than physiological IgM/IgD complexes, (ii) bearing somatically mutated rather than germline CH31 V(D)J, and (iii) lacking *in vivo* selection constraints such as self-antigen–mediated downmodulation (a hallmark of clonal anergy)^14,74^ and BCR co-receptor interactions ^75^.

To evaluate these findings under more physiological conditions, we assessed eODGT6 eODGT7, and eODGT8 immunogens—either as monomers, tetramers (low valency/avidity), and 60mer nanoparticles (np’s; high valency/avidity)—for their ability to trigger proximal BCR signals in primary splenic MF B-cells from naive CH31 UCA homozygous (VDJ^+/+^xV_κJκ_^+/+^) double KI (CH31 UCA^hom/hom^ ^dKI^) mice. This 6th–8th generation GT series was selected for two key reasons. First, despite near-identical sequence similarity, it spans a broad K_D_ range for the CH31 UCA (**Fig 1A**), enabling dissection of how association rate (k_a_) -‘on rate’- and dissociation rate (k_d_) –‘dwell time’ affect GT-mediated activation as a function of valency. Because this K_D_ range mainly reflects k_d_, this allows testing for a potential k_d_ ceiling, while the narrower range of on-rates enables verification of the minimal k_a_ (10⁴ M⁻¹s⁻¹) required by low-valency multimers to induce calcium flux in CH31 Ramos cells ^61^. Second, the eODGT8 np, represents the most characterized clinically relevant GT HIV vaccine candidate ^43^, supporting comparison to its lower-affinity predecessors. This system thus permits systematic analysis of how valency and binding kinetics influence *ex vivo* MF B-cell activation relative to *in vivo* GC priming.

**Figure 1.**
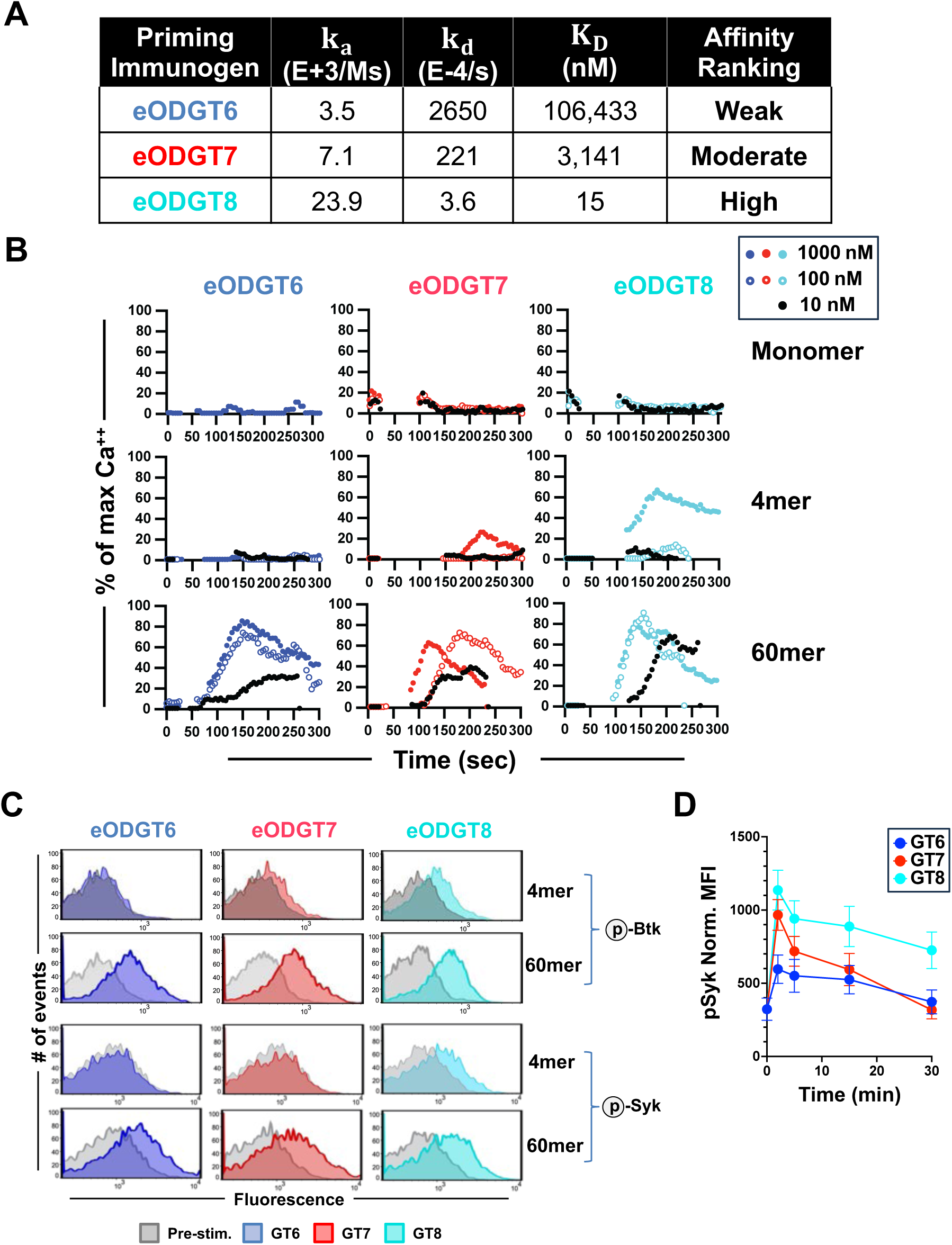
Proximal BCR signaling responses of primary CH31 UCA^hom/hom^ ^dKI^ splenic B cells to eODGT immunogens. **(A)** Binding affinities (K_D_) and kinetic parameters (k_a_ and k_d_) for eODGT6, eODGT7, and eODGT8 monomer interactions with CH31 UCA IgG, as measured by surface plasmon resonance (SPR); data are means of three independent runs. **(B)** Representative calcium flux kinetics measured by flow cytometry in CH31 UCA^hom/hom^ ^dKI^ splenic B cells stimulated with eODGT6 (low affinity), eODGT7 (intermediate affinity), or eODGT8 (high affinity) immunogens at varying concentrations and valencies. Responses are normalized to the peak induced by saturating anti-IgM F(ab′)_2_ (20 μg/ml) and are representative of two independent experiments. **(C)** Flow cytometric Mean Fluorescence Intensity (MFI) analysis of Syk (Y352) and Btk (Y223) phosphorylation at peak induction (2 min) following stimulation with either eODGT6, eODGT7, or eODGT8 (as tetramers or np’s) at saturating concentration (1 μM). Data are representative of four mice analyzed across two experiments. **(D)** Kinetics of Syk (Y352) phosphorylation following stimulation with eODGT tetramers (*top*) or np’s (*bottom*), shown as normalized MFI within live, singlet CD19^+^B220^+^ CH31 UCA^hom/hom^ ^dKI^ splenic B cells.

We first evaluated eODGT6, eODGT7, and eODGT8 immunogens, either as monomers, tetramers, or np’s, for their ability to trigger Ca²⁺ flux (a key measure of proximal B-cell signaling) in CH31 UCA^hom/hom^ ^dKI^ splenic B cells (**Fig 1B**). No immunogen induced calcium flux as monomers (**Fig 1B**, *top*), whereas eODGT8 retained notable activity as a tetramer (**Fig 1B**, *middle right*), consistent with its *kₐ* slightly exceeding the previously defined CH31 BCR activation threshold (10⁴ M⁻¹s⁻¹) for low-valency (soluble) Env trimers ^61^. Correspondingly, partial phosphorylation of proximal BCR signaling kinases Syk (Tyr532) and Btk (Tyr223), which peak by two minutes of stimulation ^61,76^, was only observed with eODGT8 tetramer stimulation (**Fig 1C**, *upper rows*). These effects were absent in WT B6 cells, confirming specificity (**Figs S1A, C**). Together, these results corroborate prior *in vitro* findings that lower-order Env immunogens require this minimal *kₐ* to activate CH31 BCRs.

In contrast, all three immunogens elicited robust Ca²⁺ flux and Syk/Btk phosphorylation when presented as high-valency, clinically relevant np’s, even with lower-affinity variants (**Figs 1B–C**, *bottom)*. This rescue of proximal signaling by increased valency (tetramer→60mer) suggests a higher CH31 BCR activation threshold in primary MF cells than in Ramos cells ^61^, warranting further investigation. We thus examined proximal BCR signaling capacity and surface density in resting splenic CH31 UCA^hom/hom^ ^dKI^ B cells (**Fig S1**), parameters associated with diminished BCR responsiveness. Intriguingly, primary CH31 UCA^hom/hom^ ^dKI^ B cells showed modest BCR downregulation (**Figs S2A,B,D**), consistent with prior self-antigen encounter ^14,62,74^, yet retained proximal BCR signaling relative to WT B-cells (**Fig S1B**). This predominantly intact signaling competence despite significantly reduced BCR density is reminiscent of a polyspecific MF B-cell subpopulation that not only preserve signaling capacity ^71^, but also have enhanced GC fitness ^23,24^, suggesting they may be particularly tractable vaccine targets. By contrast, we and others have reported more profoundly anergic bnAb precursors show only partial rescue of proximal BCR signaling and GC fitness ^37,66,69,76^, suggesting a stricter dependence on higher k_a_.

Finally, eODGT8 consistently induced the strongest proximal BCR responses across valencies. As a np, it triggered rapid, potent, and sustained Ca²⁺ flux (near 100% maximum at peak across all concentrations) and more durable Syk phosphorylation than the two earlier generation np counterparts, which decayed toward baseline by 30 minutes (**Fig 1D**). As a tetramer, eODGT8 retained ∼80% maximal Ca activity at saturating concentrations, whereas eODGT7 and eODGT6 were weak or inactive (**Fig 1B**, *middle*), and also retained substantial Btk phosphorylation (**Fig 1C**, *top*). Notably, the largest kinetic difference—eODGT8’s substantially slower rate relative to eODGT7’s—correlated with enhanced rapidity, potency, and durability of proximal signaling, indicating that higher valency and prolonged BCR engagement act cooperatively to amplify proximal signaling in CD4 mimic bnAb precursors.

### *Ex vivo* GC Differentiation of bnAb Precursors Requires Multimeric but Transient Ag Engagement

Given antigenic valency and dwell time synergize to maximally amplify k_a_-dependent proximal BCR signaling in CH31 UCA precursors (**Figs 1A–C**), we asked if these parameters become rate-limiting for distal BCR signaling programs governing clonal expansion and GC differentiation. Naïve CH31 UCA^hom/hom^ ^dKI^ splenocytes - predominantly MF B cells expressing CH31 UCA BCRs and naïve T cells - were cultured for 24h in complete media alone, or with eODGT np’s (**Fig 2A**, *top*). To enable CD4⁺ T cell help without overriding BCR signaling, splenocytes were plated in the absence of any exogenous co-stimulation, and at high but non-saturating density (5 × 10⁶/ml). 24h of stimulation was used based on our prior KI *ex vivo* studies showing peak induction of early proliferative and differentiation-associated programs in MF⁺ B cells ^76,77^.

**Figure 2.**
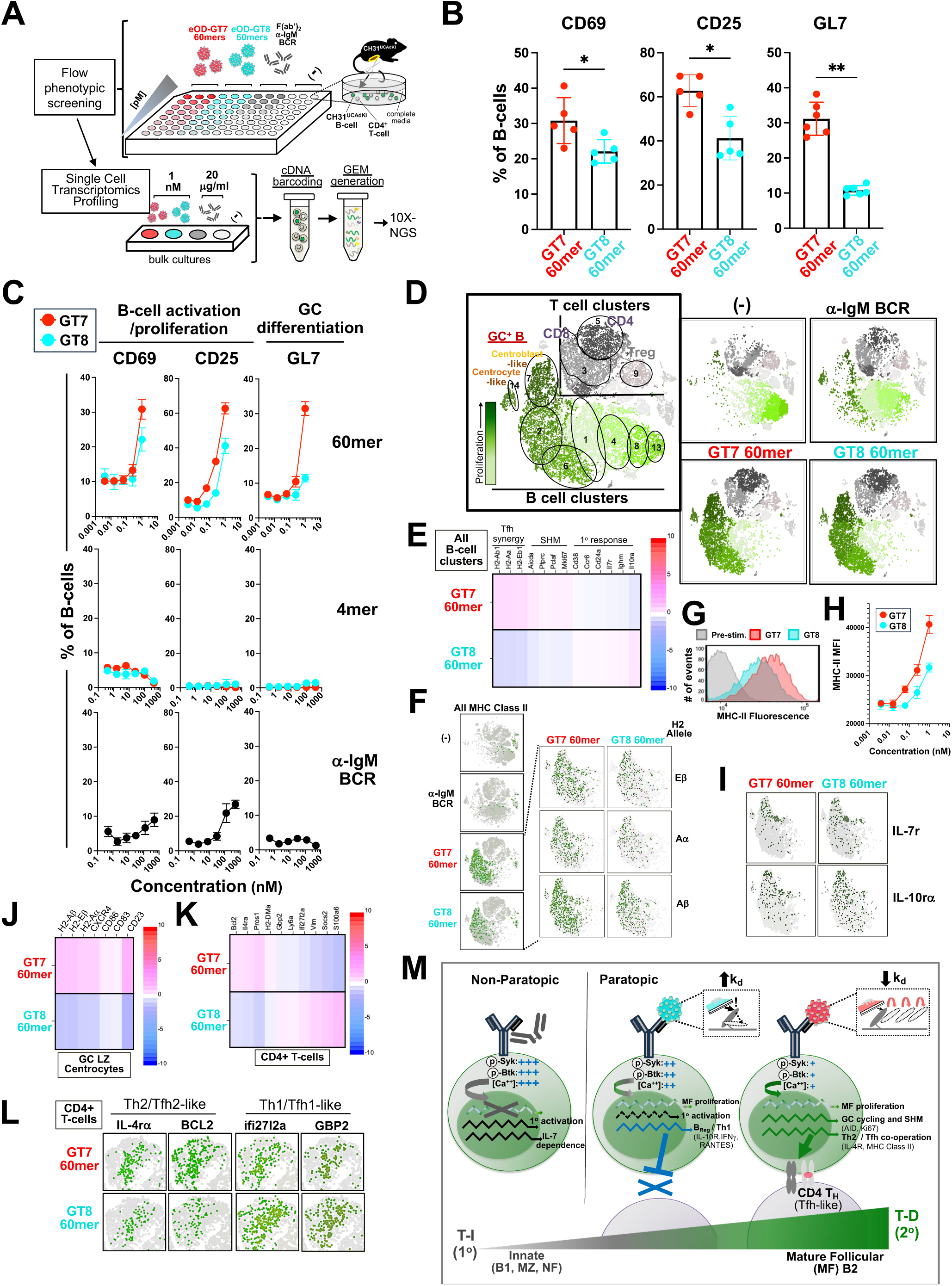
Clonal expansion and GC differentiation programs induced by eODGT7 and eODGT8 in CH31 UCA^hom/hom^ ^dKI^ splenic cultures. **(A)** Experimental workflow for ex vivo CH31 UCA^hom/hom^ ^dKI^ splenic cultures, including initial flow cytometric screening and downstream single-cell transcriptomic profiling. Cultures were stimulated in triplicate across a range of immunogen concentrations (1 pM–10 nM). Inset shows major cellular constituents. Media alone (–) served as unstimulated control. **(B)** Flow cytometric analysis of early/intermediate proliferation markers (CD69, CD25) and GC differentiation marker (GL7) on total CD19^+^B220^+^ B cells 24 h after stimulation with 1 nM eODGT7 or eODGT8 np’s. Data are expressed as percent induction over media controls; each point represents triplicate means from individual mice. Mann–Whitney U test; *p<0.05, **p<0.01. **(C)** Dose-dependent induction of CD25 or GL7 at 24 h following stimulation with eODGT6, eODGT7, or eODGT8 delivered as 60mer nanoparticles, tetramers, or anti-μ F(ab′)_2_. Values represent mean ± SEM of triplicate cultures from two mice and are representative of two independent experiments. **(D–M)** Single-cell transcriptomic profiling of CH31 UCA^hom/hom^ ^dKI^ splenic cultures. **(D)** UMAP clustering of pooled splenocytes (*n*=3) treated for 24h with eODGT7 or eODGT8 60mer nanoparticles, anti-μ F(ab′)_2_, or media alone. B-cell and T-cell clusters were defined by canonical markers; putative GC clusters were identified using curated centroblast and centrocyte gene modules. Proliferation indices highlight highest cycling in GC DZ and LZ clusters. **(E)** Heat maps comparing GC DZ-, GC LZ-, and innate/primary response–associated gene expression in B-cell clusters from eODGT7 versus eODGT8 np–treated cultures. A differential expression index is shown at right, with fold changes (−10 to +10) encoded numerically and by red (*up*) or blue (*down*) color intensity. **(F)** Single-cell expression maps of MHC class II and CD79 in B-cell clusters, shown globally and by individual MHC class II alleles in eODGT7- and eODGT8-treated cultures. **(G)** Representative flow cytometric analysis of MHC class II surface expression on total B cells after 24h stimulation, with pre-culture splenocytes shown for reference. **(H)** Quantification of MHC class II surface expression (MFI ± SEM) across a full dose range of eODGT7 or eODGT8 np’s, revealing enhanced upregulation with eODGT7, even at sub-saturating concentrations. **(I)** Single-cell maps of IL-7r and IL-10r expression in B-cell clusters, highlighting increased representation in eODGT8-treated cultures. **(J)** Differentially expressed genes in GC LZ–associated minor B-cell cluster 14, following eODGT7 versus eODGT8 np stimulation. This novel cluster was identified based on Loupe Browser differential expression using *a priori* GC LZ centrocyte gene criteria (**Fig S7**). **(K-L)** Differential gene expression (p<0.05) and single-gene maps of CD4+ T-cell cluster 3, respectively, revealing T_h_2/Tfh_2_-skewed programs with eODGT7 and T_h_1-associated signatures with eODGT8. **(M)** Model illustrating how differential eODGT7 versus eODGT8 np dissociation rates shape proximal BCR signaling vs more distal *ex vivo* BCR signals including clonal expansion, early GC differentiation, and T-cell polarization, contrasted with anti-μ–mediated non-paratopic BCR engagement.

We first quantified B cells upregulating early/late-stage proliferative surface markers CD25 or CD69 and the surface GC marker GL7, relative to media controls (**Figs 2B, C**; **Fig S3**). Strikingly, a reversal of proximal signaling hierarchies was revealed: despite higher affinity and enhanced proximal signaling by the longer-dwell time eODGT8 np’s (**Fig 1A–D**), eODGT7 np’s induced significantly greater CD25, CD69, and even more robust GL7 expression (**Fig 2B**). This advantage persisted across valency and dose, including at multiple sub-saturating (<1 nM) concentrations (**Fig 2C**, *top*), whereas tetramers were largely inactive aside from modest GL7 upregulation by saturating eODGT8 tetramer (**Fig 2C**, *middle*). In contrast, non-paratopic anti-Ig signals (via Framework; FRW and/or Cμ region interactions) weakly induced CD25/CD69 and failed to induce GL7, even suppressing it at super-saturating doses (**Fig. 2C**, *bottom*). Thus, in contrast to proximal signaling, moderate-affinity, faster-dissociating eODGT7 np’s more effectively promote early GC differentiation than higher-affinity, slower-dissociating eODGT8 np’s.

To further define these distal programs, we performed single-cell RNA-seq on CH31 UCA^hom/hom^ ^dKI^ splenic cultures treated for 24 h with eODGT7 or eODGT8 np’s, anti-IgM, or media alone (>20,000 cells per condition; **Fig 2A**). Unbiased clustering identified major T cell (CD3^+^CD4^+^/CD8^+^) and B cell (CD79α/μ^+^) populations (**Fig 2D**). Within B-cells, eODGT np’s selectively enriched clusters 2, 6, and 7, whereas anti-IgM preferentially enriched clusters 1, 4, and 8; cluster 13 was unique to media (**Fig 2D**). Notably, eODGT np-expanded B-cell clusters exhibited robust proliferative programs, including gene expression involved in DNA replication and repair (*PCNA, MCM3/4, Hells*), cell-cycle regulation (*Siva1*), and nucleotide metabolism (*Dut, Dtymk*), forming a continuum from clusters 2 to 7. Even more remarkably, cluster 7 displayed a GC dark zone (DZ) centroblast-like transcriptional profile, with additional expression of SHM-associated DNA repair/replication genes like *Top2a and Pclaf* (**Figs S4, S5**). In contrast, anti-IgM–treated B cells showed minimal proliferative pathway induction and instead upregulated markers associated with quiescent naïve or memory B-cell states (*Ly6d, CD38, CD22, CD55, Bank1*) (**Fig S5**). Thus, paratopic engagement of CH31 UCA^hom/hom^ ^dKI^ BCRs by eODGT np’s efficiently drives highly proliferative B-cell states (consistent with FM clonal expansion), in contrast to weak, less-specific, activation induced by non-paratopic anti-IgM stimulation (**Figs 2B, C**).

Building on this shared capacity of eODGT7 and eODGT8 np’s to drive proliferative clonal expansion, we next determined if and how their transcriptional programs differed. Despite highly similar global clustering (**Fig 2D**), differential gene expression analysis revealed clear divergence between eODGT7- and eODGT8-induced programs (**Figs 2E,F,I,J**). eODGT7 np–stimulated CH31UCA^hom/homdKI^ B cells preferentially upregulated genes associated with GC proliferation and function (*MKi67, Pclaf, AID*), together with Tfh pathway (Th2-synergizing) genes, including IL4Rα and MHC class II across all H2 alleles (**Figs 2E,F**). Enhanced MHC class II expression was confirmed at the surface protein level, including under sub-saturating np conditions (**Figs 2G,H**). In contrast, eODGT8 favored expression of genes linked to primary/innate activation (*IgM, CD38, IL7r*), premature GC exit (*CCR6, HSA*), and Th1 skewing (Tfh-dampening) related genes including IL10Rα (**Figs 2E,I**). These opposing transcriptional programs were consistent across individual clusters, including differential MHC class II, core GC proliferation genes, and IL7r/IL10Rα expression (**Fig S6**).

Subclustering further resolved these distinctions (**Figs S7, S8**). A minor cluster adjacent to major cluster 7 (**Figs 2D, S7a**) displayed a GC light zone (LZ), centrocyte-like profile (*CD83, CD86, CXCR4, CD23*, high *MHC class II*; **Figs 2J, S7b**), whereas cluster 7 itself exhibited a GC DZ–like signature marked by elevated proliferation and SHM-associated genes (*Pclaf, PCNA, Top2a*, *AID*) (**Figs 2D, S7b**). Notably, eODGT7-treated cultures were significantly enriched for both these putative DZ and (especially) LZ GC subsets, reflected by differential expression of signature genes (**Figs sS7b**) and increased representation of clusters 7 and 14 (**Fig S7c**). Conversely, eODGT8 selectively expanded high IL10Rα-expressing B cells across proliferative clusters (1, 2, 6, 7, 14; **Figs 2I** *lower*, **S8a**) consistent with a putative B_Reg_ (B10) phenotype (**Fig S8b**) ^78,79^, including increased CCL5 and interferon-regulated genes (CXCL16,CD9, CCR2) (**Fig 8b**) associated with suppressing adaptive T_h_2/Tfh responses ^80,81^.

Consistent with these B-cell–intrinsic programs, CD4^+^ T-cells from eODGT7-treated cultures preferentially adopted a Tfh2-like profile with enhanced survival and GC-supportive genes (*BCL2, IL4r, H2-DMa*), whereas eODGT8 treatment promoted interferon-associated, Th1-skewed programs (*GBP2, IFI27l2a, Ly6a*) with limited global induction (**Figs 2K,L, S9A**). CD4^+^ Treg and CD8^+^ T-cell populations were not significantly altered (**Figs S9A; S11**). Also of further interest, Anti-IgM uniquely upregulated IL7r in CD4+ T cells (***Fig S9B***), in line with its non-paratopic interactions with CH31 UCA^hom/hom^ ^dKI^ B-cells.

Collectively, these distal (24h) *ex vivo* BCR signaling readouts—surface phenotyping and genome-wide transcriptional profiling—indicate that: (i) antigenic valency sets a threshold for proliferation and GC entry (ii) in contrast to proximal signaling, slower dissociation (k_d_) signals delivered by high affinity eODGT8 are supra-optimal, diverting away from GC differentiation, proliferation/SHM, T_h_2/Tfh synergy, and/or GC retention programs relative to eODGT7; and (iii) non-paratopic anti-IgM stimulation induces a weak, IL7-dependent, T-independent/innate-like response, associated with less mature B-cell populations, while eODGT8 produces an intermediate phenotype between these and optimal T-dependent responses. More broadly, productive engagement of bnAb precursors in GCs requires GT interactions at sufficient association rate at high valency, while avoiding excessively slow dissociation that activate primary response programs. As we model using a ‘traffic flow’ analogy in **Fig 2M**, k_a_ governs entry, valency sustains progression, and k_d_ functions as braking—where excessive braking (sustained engagement) promotes premature GC exit, whereas more rapid dissociations (allowing multiple re-associations/‘brake pumping’, as with eODGT7) support continued GC retention, proliferation, and differentiation.

### Developing a CH31 transfer model for testing *in vivo* GC priming

While our above-described *ex vivo* culture platform demonstrates that faster-dissociating eODGT7 np’s more effectively activate GC-associated transcriptional and Tfh2 programs than slower-dissociating eODGT8 np’s, these findings derive from a splenic culture system maximizing testing rapidity rather than physiologic complexity. This system contains near-homogeneous B-cell specificities with minimal clonal competition and less organized T helper (T_H_) cell signals, which may dampen rather than amplify eODGT7 selectivity *ex vivo*, relative to physiologic conditions where antigen availability is limiting and diverse off-target B-cell clones compete for alternative eOD epitopes. Conversely, *in vivo* factors could attenuate kinetic advantages, including low bnAb precursor frequency ^53,55^, altered basal BCR signaling thresholds due to self-reactivity (**Fig 2A**), and stronger, spatially organized Tfh-derived GC co-stimulation that may partially override intrinsic BCR signaling differences observed *ex vivo*.

Thus, to test whether *ex vivo* kinetic differences predict *in vivo* priming and to define how binding kinetics interact with precursor frequency and clonal competition, we developed an adoptive transfer–based CH31 precursor KI model with precisely controlled precursor abundance (**Fig 3**). CH31 UCA^hom/hom^ ^dKI^→WT chimeras were generated via retro-orbital transfer of negatively purified splenic B cells from CH31 UCA dKI donors into congenic wild-type recipients (**Fig 3A**). Model performance was validated by titrating donor B-cell inputs over a 3–10-fold range (10²–3×10⁷ cells) and quantifying donor recovery 24h post-transfer (**Fig 3B).** Across three independent experiments, cohorts receiving ∼10⁷–2×10⁴ input cells displayed highly reproducible input/output ratios (∼750:1), defining a near-linear reconstitution window corresponding to ∼10⁻³–10⁻⁶ CH31 precursor frequencies in recipient spleens. This range spans immunodominant precursor frequencies typical of murine hapten or protein immunizations ^82–85^, intermediate frequencies associated with conventional antiviral nAb responses ^16,86,87^, and estimated frequencies of CD4-mimic bnAb precursors in humans ^34,88^, which exist in most humans at higher frequencies than most other bnAb precursor classes ^89^. The lowest reproducibly detectable frequency (∼10⁻⁶) corresponds to absolute precursor numbers in mice (∼10–100 cells) consistent with ultra-rare bnAb clonotypes in humans (<10⁻⁹), after accounting for differences in peripheral immune compartment size (**Fig 3B**). Together, these data establish an *in vivo* platform in which CH31 UCA^hom/hom^ ^dKI^ precursor frequencies can be reproducibly tuned across physiologically relevant immunodominance tiers, enabling direct assessment of how GT immunogens with distinct binding kinetics can prime GC fitness in a prototype CD4-mimic bnAb lineage.

**Figure 3.**
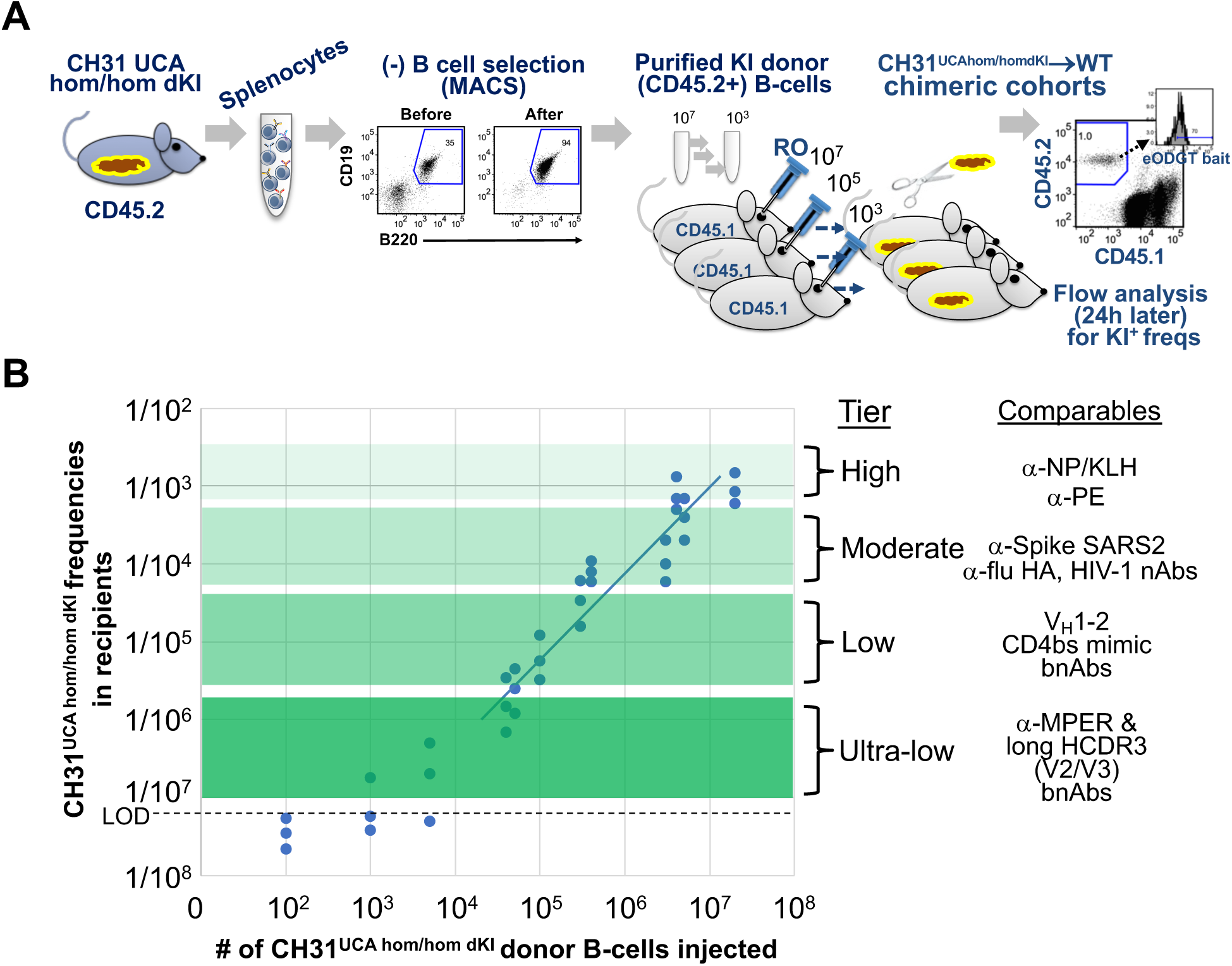
Development of an *in vivo* CH31 precursor B-cell transfer model. **(A)** Schematic of transfer process, titration experiments, and downstream repertoire analyses. **(B)** Determination of donor input numbers required for reproducible reconstitution of B6.SJL CD45.1+ recipients with CD45.2+ CH31 UCA^hom/hom^ ^dKI^ B cells. Shown are input–output relationships between injected donor cells and recovered CH31 precursors within the splenic B-cell compartment 24 h post-transfer, quantified by flow cytometry using CD45.2 and eODGT bait discrimination. Each point represents an individual chimera; data are pooled from three independent experiments (≥3 mice/group). The level of detection (LOD) reflects limits imposed by total splenic B-cell numbers and flow cytometric sensitivity. Shaded regions denote defined precursor frequency tiers: high (<1:10^3^), moderate (∼1:10^4^), low (∼1:10^5^), and ultra-low (∼1:10^6^), with comparables at each tier indicating representative germline precursors for canonical murine neo-antigen responses and human viral vaccine targets.

### Transient Immunogen–BCR interactions drive GC Fitness across all bnAb precursor frequencies

Given the importance of precursor frequency for informing the eOD np clinical platform, we used the CH31 precursor transfer model to assess how this parameter influences the ability of eODGT7 and eODGT8 np’s to recruit CH31 precursors into GCs (**Fig 4A**). CH31 UCA^hom/hom^ ^dKI^→WT chimeras were generated with 10-fold–decreasing CH31 precursor frequencies spanning the reproducible 10⁻³–10⁻⁶ range, corresponding to four immunodominance tiers (**Fig 3B**). Following an 8-day prime, eODGT7 consistently recruited a significantly larger fraction of CH31 precursors into GCs than eODGT8 (**Fig 4B**).

**Figure 4.**
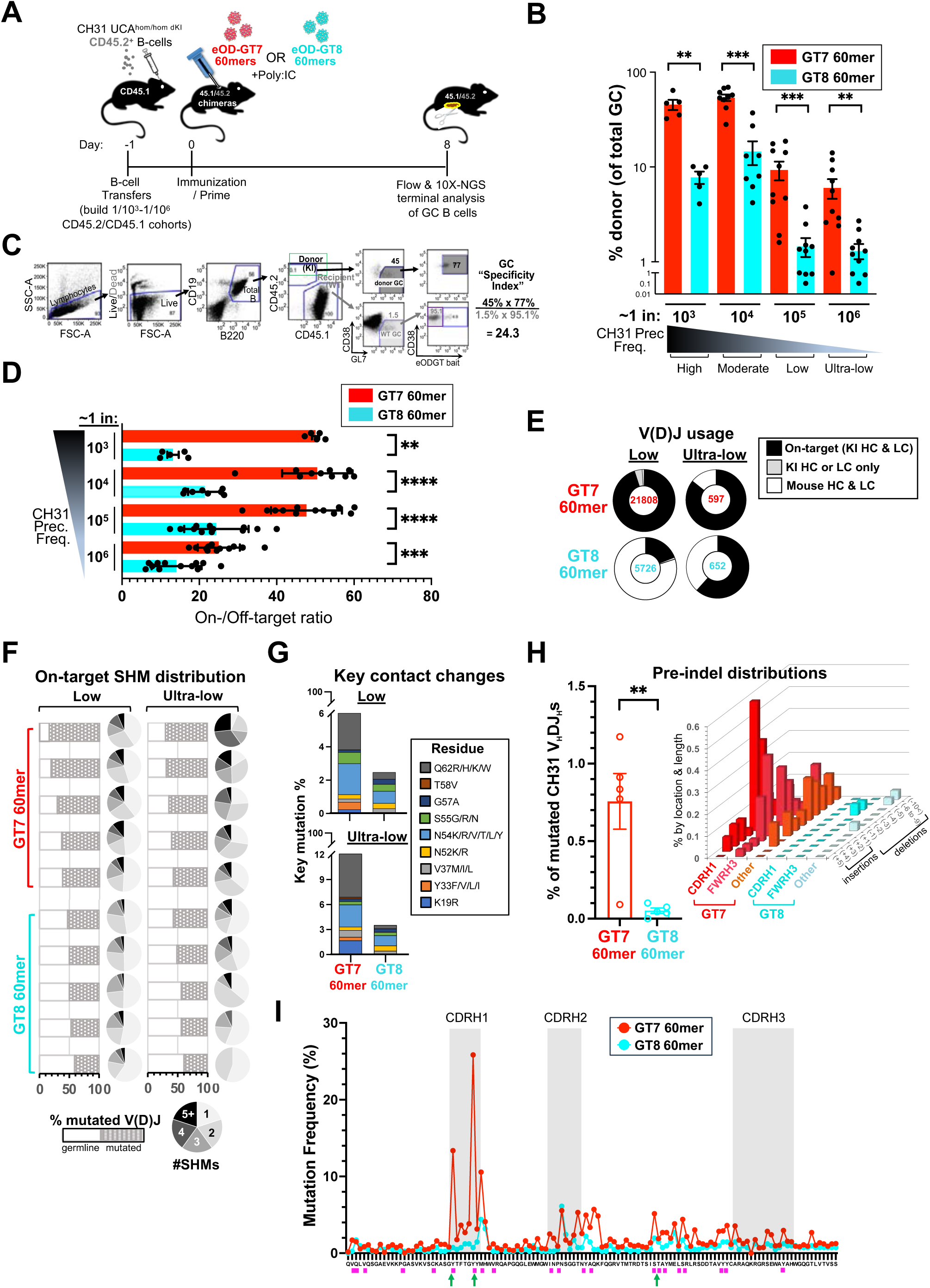
Effect of precursor frequency on eODGT7 versus eODGT8 np GC recruitment and SHM. **(A)** CH31 precursor adoptive transfer and priming schema. B6.SJL CD45.1+ mice were reconstituted with graded numbers of CD45.2+ CH31 UCA^hom/hom^ ^dKI^ B cells (Fig 3; Methods) and primed for 8d with eODGT7 or eODGT8 np’s. **(B-D)** Flow cytometric analysis of GC recruitment in primed CH31 UCA^hom/hom^ ^dKI^→WT chimeras. **(B)** GC occupancy, defined as the percentage of GC B cells (B220+CD19+CD38–GL7+) that are donor-derived (CD45.2+) and ‘on-target’ (eODGT bait+, KI HC+LC). **(C)** Representative gating strategy for calculation of the GC ‘specificity index’ ie, on-target donor GC B cells divided by ‘off-target’ (CD45.1+eODGT-) recipient GC B cells. **(D)** GC recruitment specificity index (on/off-target ratio) following eODGT7 or eODGT8 np priming. Each point represents one primed chimera. **(E–I)** 10x paired HC/LC sequencing and SHM analysis of individual GC B cells from primed chimeras reconstituted at low (∼1:10^5^) or ultra-low (∼1:10^6^) precursor frequencies, corresponding to physiological human estimates and the lower limit of reproducible reconstitution, respectively (**Fig 3B**)^34,64^. **(E)** HC/LC pairing status of single-sorted donor GC B cells, with *bona fide* (V(D)J sequence verified) on-target clones indicated. Pies show HC/LC pairing composition of donor GC B cells; black slices mark on-target CH31 UCA pairs and center values denote unique cell counts. **(F)** Total SHM distributions among on-target CH31 clones, with bar graphs showing the fraction of on-target CH31 clones that remain germline versus those acquiring ≥1 aa substitution, and accompanying pie charts stratifying mutated clones by aa substitution number. **(G)** Frequency of VRC01/CH31-class key V_H_ contact residue substitutions among mutated on-target clones^19^. **(H)** Frequency and distribution of pre-indel events in CH31 HC rearrangements, stratified by HC region and indel length^19,115^. **(I)** Positional distribution of CH31 HC aa mutations across on-target pairs, with AID hotspot motifs (WRC/GYW and WGCW; W =A/T, R=A/G, Y=C/T) indicated by blocks and previously reported insertion sites denoted by arrows ^18,19,116^. Data in panels (F) and (H) represent pooled 10x Ig-seq single cells from chimeras over two independent experiments (five pools; 18 mice per 1/10^5^-reconstituted group; 6 mice per 1/10^6^-reconstituted group, *n*=48 total). Data in panels (G) to (I) comprise chimeras reconstituted at physiological (1/105) precursor frequencies. Statistical comparisons used Mann–Whitney U tests. *p<0.05, **p<0.01, ***p<0.001, ****p<0.0001; ns, not significant.

To quantify GC fitness and specificity, we developed a flow-based GC bnAb specificity index measuring the ratio of on-target CH31 precursors to ‘off-target’ (non-bnAb) competitors (**Fig 4C**). By this metric, eODGT7 demonstrated markedly higher CH31-specific GC recruitment than eODGT8 (**Fig 4D**), an advantage that was maintained across all precursor frequencies, including the estimated physiologic mean (∼1/100,000). At ultra-low precursor frequencies, overall clonal expansion did not differ significantly between eODGT7- and eODGT8-primed animals (**Fig S12**), reflecting rate-limiting conditions due to precursor number, and consistent with *ex vivo* transcriptional profiling showing comparable proliferative programs (**Fig 2D**) when BCR and CD4 T-cell interactions are limiting. However, differences in GC/Tfh fitness programs remained evident, indicating that intrinsic BCR signaling kinetics selectively shape GC recruitment and differentiation rather than proliferation. Consistent with this, DZ/LZ ratios also remained unchanged (**Fig S13**), supporting the conclusion that sufficient expansion occurs, GC differentiation programs are intrinsically driven by their BCR sensing kinetics.

To determine whether enhanced GC recruitment by eODGT7 np’s enhances the mutational landscape available for affinity maturation, we performed high-throughput paired heavy- and light-chain sequencing on single GC⁺ B cells sorted from eODGT7- or eODGT8-primed recipients, seeded with low/∼physiologic (10⁻⁵) or ultra-low (10⁻⁶) CH31 precursor frequencies **Fig 3B**). Immunogenetic verification was required due to potential LC editing secondary rearrangements *in vivo*^19^. Consistent with flow-cytometry (**Fig 4B-D**), a significantly higher fraction of GC⁺ B cells from eODGT7-primed animals expressed verified CH31 UCA V(D)J rearrangements (**Fig 4E**). Among these on-target clones, eODGT7 priming resulted in a higher proportion of mutated sequences and then within these, greater mutational frequencies were observed compared to eODGT8, with differences amplified at ultra-low precursor frequencies (**Figs 4F**). Notably, eODGT7 np priming preferentially induced single-amino-acid substitutions required for CD4-mimic bnAb maturation (**Fig 4G**), and remarkably, multi-residue HCDR1 and FRW3 indels (**Fig 4H**), which are crucial for broad Env recognition ^17–19^. These indels were almost exclusively observed in eODGT7-primed clones and were predominantly large deletions, consistent with proposed “pre-indel” intermediates in CH31–CH34 lineage evolution ^17,18^. Supporting this, eODGT7-primed clones exhibited increased SHM at, and flanking, AID hotspot–rich regions within HCDR1 and FRW3 (**Fig 4I**), recapitulating patterns previously identified during CH31-34 bnAb evolution in infection^18^ and vaccination^19^.

Together, these *in vivo* data using the CH31 transfer model validate our *ex vivo* distal findings of an intrinsic k_d_-delimited ceiling in the eODGT platform’s ability to prime GC bnAb responses, here showing this occurs regardless of bnAb precursor frequency.

### *In vivo*, GT immunogen valency only partially modulates the k_a_/k_d_-defined GC priming range

We next tested whether the second major prediction of the *ex vivo* CH31 precursor culture system — that antigenic valency modulates the k_a_/k_d_-defined GC fitness window — holds *in vivo*. *Ex vivo*, increased valency partially overcame a minimal association rate required for GC differentiation, whereas reduced valency minimally mitigated excessively long dwell times (**Fig 2C**), although these studies were performed under rate-limiting conditions where extrinsic constraints (ie sub-optimal ag, clonal competition, or T cell help) may have influenced these conclusions. Thus, to formally test this *in vivo*, we used the CH31 precursor transfer model to seed WT recipients at the mean physiologic precursor frequency (∼1/100,000) (**Fig 5A**) and immunized with monovalent, low-valency (4mer), or high-valency (60mer np) forms of eODGT6, eODGT7, and eODGT8, spanning the full kinetic range. GC responses were assessed at d8 (peak GC occupancy) ^53,63,90^ and d16 (SHM plateau) ^53–55,91^.

**Figure 5.**
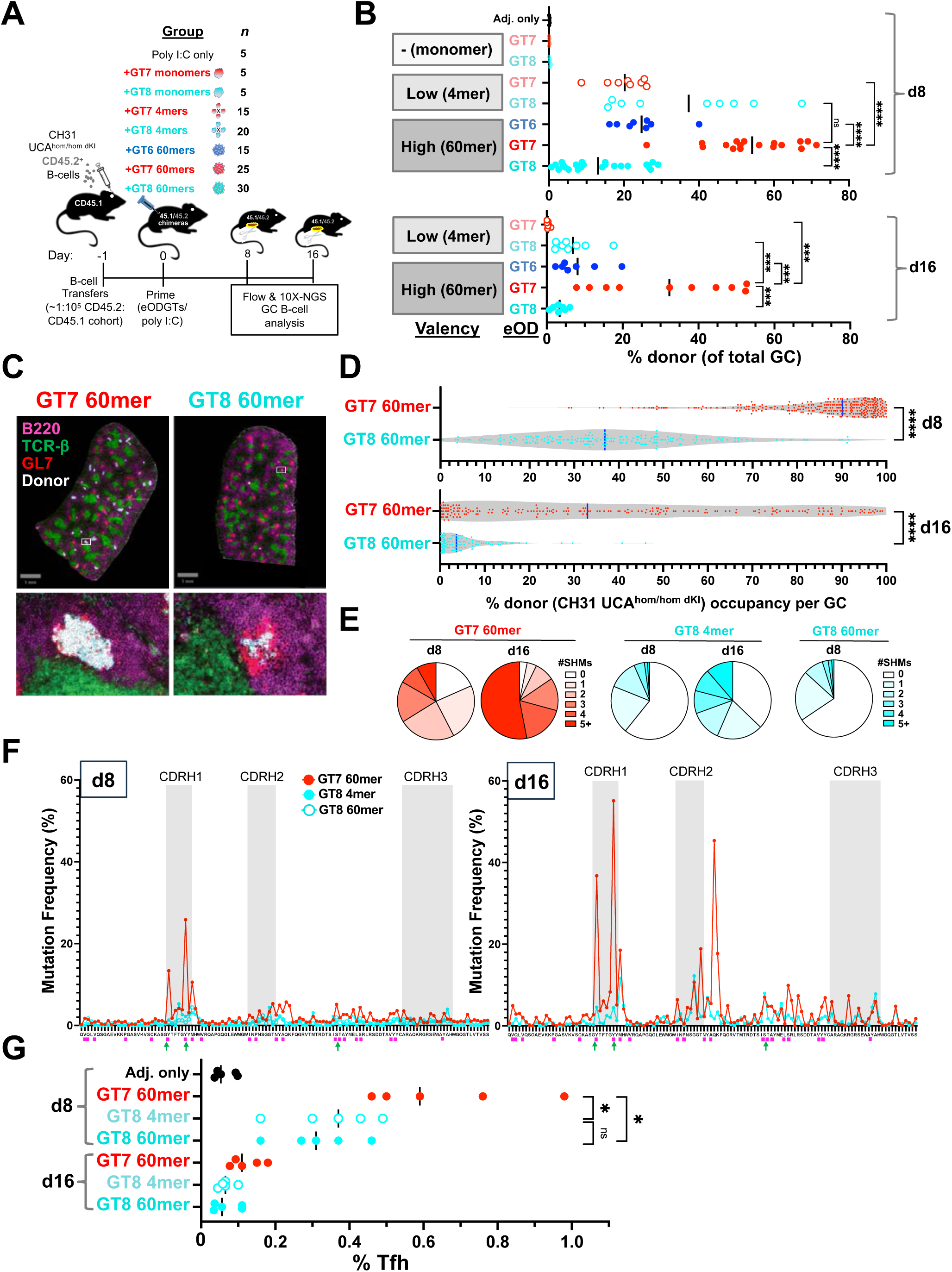
Effects of eODGT6–8 platform’s k_a_/k_d_ and valency on magnitude and persistence of CH31 precursor GC priming and V(D)J SHM. **(A)** Schematic of the CH31 precursor transfer and priming strategy. B6.SJL CD45.1 recipients received purified CD45.2⁺ CH31 UCA^hom/hom^ ^dKI^ B cells and, after 24h, resulting CH31 UCA^hom/hom^ ^dKI^→WT chimeras (reconstituted at 1:10^5^ precursor frequency) were primed for 8 or 16 days with eOD monomers, low-valency tetramers, or high-valency np’s (poly I:C–formulated), or poly I:C alone, prior to GC B cell recovery for flow cytometric phenotyping and 10x paired Ig seq (*n* per group indicated). **(B)** Flow cytometric quantification of donor-derived, on-target GC B cells (live singlet B220⁺CD19⁺CD38⁻GL7⁺CD45.2⁺eODGT bait⁺) shown as the fraction of total GC B cells at day 8 (*top*) or day 16 (*bottom*). Each dot represents one primed CH31 UCA^hom/hom^ ^dKI^ chimera. Significance relative to eODGT7 np was determined by Mann–Whitney U tests. *p<0.05; **p<0.01; ***p<0.001; ****p<0.0001. **(C–D)** Immunofluorescence validation of enhanced CH31 precursor recruitment and GC persistence following eODGT7 np priming. **(C)** Representative spleen cryosection images from chimeras primed for 16 days with eODGT7 or eODGT8 np’s; boxed regions indicate GC closeups. Note the persistently high fraction of donor CH31 UCA^hom/hom^ ^dKI^ (CD45.2⁺) B cells in individual GCs across the spleen (top) of an eODGT7 np–immunized animal at this later time point. **(D)** Quantification of donor CH31 GC occupancy (CD45.2⁺ area within GL7⁺ GCs) at days 8 and 16, shown as violin plots with medians (blue bars). Each dot represents one GC. Data are pooled from three mice per group. Total GCs analyzed were 698/465 (eODGT7, d8/d16) and 505/325 (eODGT8, d8/d16). Mann–Whitney test; ****p<0.0001. **(E-F)** SHM magnitude, positional distribution in, and kinetics of, CH31-derived V(D)J rearrangements following eODGT7 np priming, compared with eODGT8 np or eODGT8 tetramer priming. GC B cells were isolated by flow sorting and analyzed by paired 10x Ig-seq. **(E)** Pie charts show SHM distributions among *bona fide* on-target CH31 UCA HC/LC pairs, with slices indicating total aa substitutions. ≥1000 unique, non-oligoclonal sequences from ≥5 pooled mice per group were analyzed. Data from eODGT8 np–primed mice are omitted due to insufficient recovered precursors. **(F)** Frequency of CH31 UCA HC aa mutations by residue position among all on-target pairs. AID hotspots (WRC/GYW and WGCW) are indicated by pink blocks. Previously reported insertion sites from bnAb lineage retracement ^18,116^ or vaccine-induced maturation ^19^ are marked by arrows. **(G)** T follicular helper (T_fh_) cell responses in chimeras primed for 8 or 16 days with eODGT7 or eODGT8 np’s, or eODGT8 tetramers. Shown is the percentage of live, singlet splenocytes that were T_fh_ (CD4⁺CD44⁺CD62L⁻PD1⁺CXCR5⁺CD25⁻CD127⁺) at peak GC occupancy (d8) or peak SHM (d16). Gating strategy is shown in Fig S16.

At d8, faster-dissociating eODGT7 np’s consistently recruited the highest fraction of CH31 precursors into GCs among all immunogens tested, including eODGT8 np’s (**Fig 5B**, *top*), consistent with superior GC cycling dynamics. Reduced valency partially improved GC entry, as lower-valency eODGT8 4mers showed intermediate recruitment. However, by d16, eODGT7 np–primed animals retained significantly more CH31 precursors than all other groups, including eODGT8 4mers (**Fig 5B**, *bottom*), demonstrating that only eODGT7 np priming achieves sustained GC occupancy (the priming ‘sweet spot’). Histology confirmed greater GC occupancy by CH31 donor cells at both time points following eODGT7 np priming (**Figs 5C,D**).

To further examine if reduced valency partially alleviates the k_d_-defined GC fitness ceiling, we compared SHM in CH31 UCA rearrangements following priming with eODGT7 np’s eODGT8 np’s, or eODGT8 4mers (**Figs 5E,F**). Due to low precursor recovery in both eODGT8 groups at d16, analyses focused on the earlier (d8) sampling time. At this time point, eODGT8 4mers induced only marginally greater SHM than eODGT8 np’s and remained markedly inferior to eODGT7 np’s in overall mutation frequencies and acquisition of functionally key HCDR1 and FRW3 amino acid replacements (**Figs 5E,F,S15**). Consistent with superior GC cycling, eODGT7 60mers also elicited significantly stronger T_fh_ responses at d8 than either eODGT8 immunogen, including the intermediate-performing 4mer, without altering Treg frequencies or T_fh_/T_reg_ ratios (**Figs 5G, S16**). Thus, while reduced valency of slow-dissociating eODGT8 partially improves GC entry (consistent with partial rescue of early GC differentiation in eODGT8-stimulated CH31 cultures) it ultimately fails to overcome the intrinsic requirement for transient (rather than prolonged) BCR–antigen interactions to sustain GC fitness, SHM accumulation, and T_fh_ support.

At the opposite boundary (the priming “floor”), valency likewise cannot compensate for insufficient association kinetics. High-valency eODGT6 np’s exhibited reduced CH31 precursor GC recruitment and retention (**Fig 5B**), indicating that increasing valency cannot compensate for a sub-threshold k_a_. Similarly, priming with lower-valency eODGT7 4mers showed impaired CH31 precursor recruitment to GC relative to eODGT7 np’s, indicating that immunogens right near the k_a_ threshold require higher valency to achieve robust entry.

Collectively, our results of GC priming with all forms of eODG6, -7, or -8 in CH31 UCA^hom/hom^ ^dKI^→WT chimeras demonstrate a constrained GC priming window *in vivo*: valency modulates but cannot override intrinsic kinetic limits. It partially tunes the k_a_-defined priming floor but fails to rescue GC occupancy or retention below it nor does it alleviate the k_d_-defined ceiling. Strikingly, and directly relevant for reducing effort and resources involved in empirical vaccine testing, all valency-dependent effects on CH31 precursor GC fitness closely mirror the early GC differentiation signals predicted by the 24h ex vivo assay across the same GT immunogens (**Figs 2, S4–S11**), establishing this system as a predictive framework for in vivo bnAb precursor priming. A model that collectively incorporates these novel concepts for optimal GC activation is shown in **Fig 6**.

**Figure 6.**
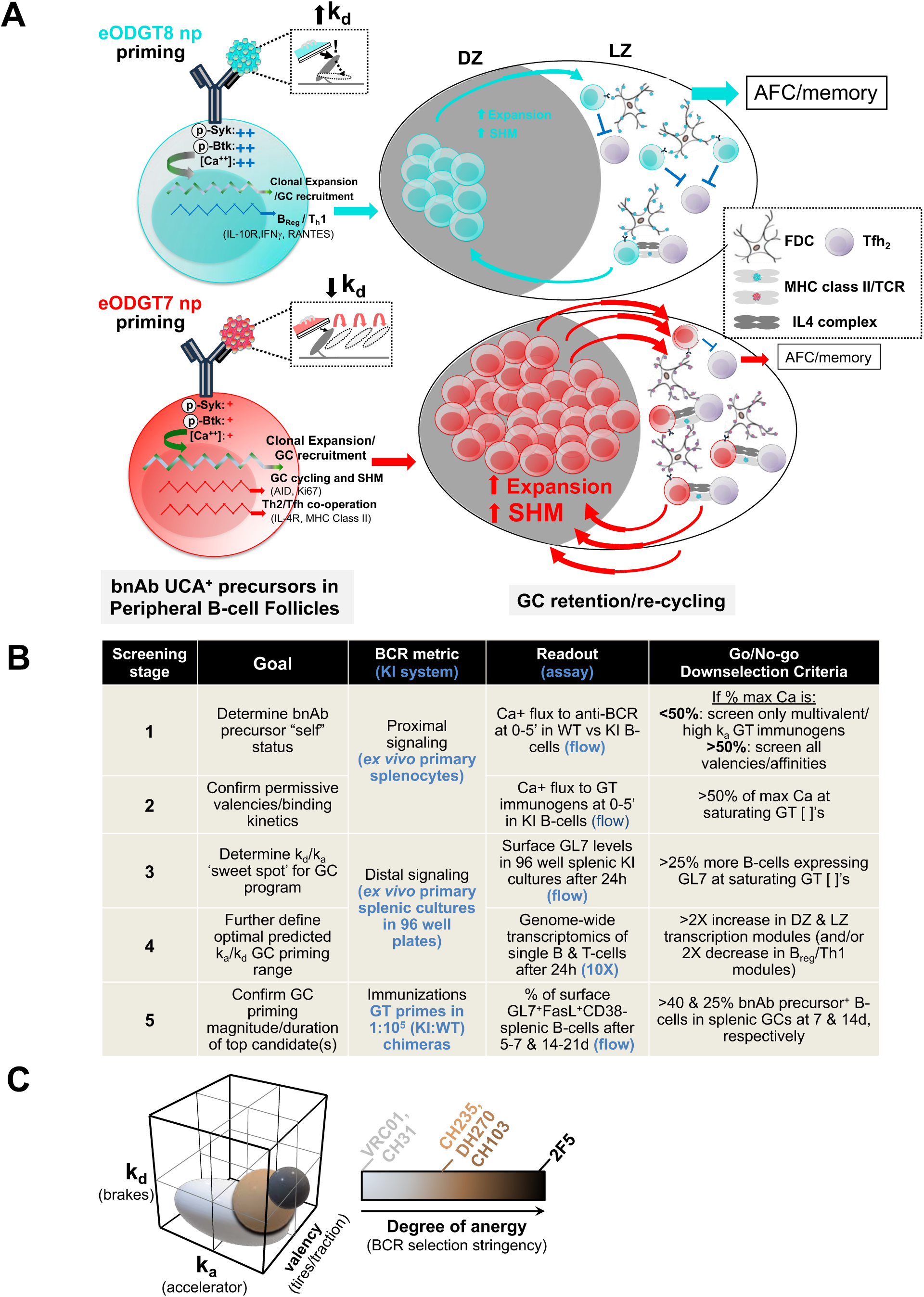
A predictive framework linking GT immunogen binding kinetics and valency to early GC programs *ex vivo* and GC fitness/SHM priming *in vivo*. **(A)** Global traffic flow model describing how bnAb precursor BCR sensing of GT immunogen dwell time (k_d_) intrinsically regulates GC DZ/LZ transcriptional programs within peripheral follicles, collectively increasing bnAb precursor recruitment, cycling, and retention in GCs. An optimal, intermediate-range k_d_ maximizes GC fitness—from GC gene module induction to recruitment, to DZ/LZ recycling—analogous to light/controlled, repetitive braking that keeps a car moving in heavy rush-hour traffic. In contrast, excessively slow k_d_ resembles hard and sustained ‘emergency’ braking, halting progress and diverting responses toward alternate outcomes (e.g. short-lived memory or AFC differentiation above ceiling). FDC, follicular dendritic cell; AFC, antibody-forming cell/plasmacyte. **(B)** Proposed workflow integrating BCR binding metrics, experimental readouts, and go/no-go criteria to enable more efficient preclinical ranking and down-selection of GT immunogen candidates in KI models. This framework prioritizes candidates capable of priming lineage-specific bnAb precursors into GCs and increasing the likelihood of improbable SHM events, while limiting *in vivo* studies largely to confirmation, thereby accelerating progression to clinical testing. **(C)** 3D schematic illustrating how increasing stringency of peripheral negative selection (“self” status) progressively narrows the optimal GC priming range (“sweet spot”). Required GT immunogen dissociation rate (k_d_) is shown on the y-axis (braking force), while association rate (k_a_; accelerator) and valency (traction) are shown on x- and z-axes, respectively. Increasing negative selection requires greater acceleration (higher k_a_) when traction (higher valency) is no longer limiting, in order to sustain GC priming under more restrictive conditions. Colored spheres denote predicted GC priming sweet spots for bnAb precursors under varying degrees of host control, from minimal (lighter) to stringent (darker).

## DISCUSSION

A preventive HIV vaccine will likely need to activate rare and/or disfavored precursor B cells and guide their maturation into bnAbs targeting conserved sites on trimeric Env^3,7,15,92^. The CD4BS represents a major target, and germline-targeting (GT) vaccine design work has thus focused on V_H_-restricted (‘CD4 mimic’) bnAbs such as the VRC01/CH31 class^41,93–96^. However, eliciting broadly neutralizing sera in humans remains challenging, in part because the biophysical requirements of GT immunogens to support robust precursor clonal expansion and GC fitness during priming remain poorly defined. Using the CH31 UCA dKI model^19^, we systematically examined how varying binding kinetics and valency across the eOD-GT platform shapes early *ex vivo* GC transcriptional programming and *in vivo* GC recruitment, cycling, and retention. These studies define an optimal kinetic window for GC priming at physiological precursor frequencies in which sufficiently high association rates together with intermediate dissociation rates promote sustained GC fitness, whereas excessive dwell time or subthreshold association rates impair GC persistence.

These findings have several profound implications for both HIV vaccine design and basic B-cell immunology. First, they confirm the importance of sufficiently high association rates^60,61^ and antigen valency^28,65,66,69^ for GT-initiated bnAb lineage development while establishing binding kinetics - including a previously unrecognized dwell-time ceiling - as predictive determinants of bnAb precursor GC priming. They also challenge the prevailing (though not unanimous;^55^) view that higher affinity is optimal at physiological precursor frequencies^52–54^. Instead, our results offer an alternative interpretation of a pre-clinical KI transfer study^53^ that was seminal in advancing the eOD-GT8 np candidate into phase I HIV vaccine trials: its success may have reflected a sufficiently high k_a_ compared with earlier eOD-GT generations (eOD-GT2 and eOD-GT5), which fell below the defined kinetic threshold^61^. More broadly, these findings address a longstanding question in immunology - how B cells interpret the biophysical parameters of antigen-BCR engagement to regulate GC fate decisions-and provide a framework for more efficient selection and optimization of GT immunogens targeting HIV and other pathogens requiring exceptional antibody breadth and durability.

Disentangling association rate, dwell time, and valency from equilibrium affinity further revealed-within the eOD-GT6/7/8 monomer, tetramer, and 60-mer series-the unique capacity of eOD-GT7 np’s to both initiate and sustain GC responses at physiological frequencies of CH31–CH34 lineage precursors. These properties indicate that the combined valency and binding kinetics of GT7 align with a GC-priming “sweet spot” for this bnAb lineage. Integrating temporally linked readouts—from proximal signaling in untouched primary B cells to *in vivo* GC recruitment and SHM—we propose a clonotype-specific “traffic flow” kinetic model (**Fig 6A**) in which k_a_/k_d_-dependent paratope sensing calibrates the balance between Tfh-driven GC programs and primary effector outputs (unmutated AFC and/or short-lived T-dependent memory). Although both eOD-GT7 and eOD-GT8 np’s engage the CH31 paratope, their distinct dissociation rates encode divergent fates: GT7 (transient BCR engagement) drives a fully T_h_2-skewed, GC-competent program with coordinated DZ/LZ cycling, whereas GT8 (sustained engagement) supports follicular proliferation but limited GC recruitment, representing an intermediate paratopic state between innate-like activation and fully GC-skewed signaling. By contrast, general anti-Ig stimulation—mediated via non-paratopic framework and/or constant-region interactions—preferentially activates IgM^hi^ innate subsets (marginal zone, transitional/newly formed, B1). Notably, proximal signaling events mirror 24-hour GC and SHM-permissive transcriptional programs, indicating that early kinetic cues preconfigure downstream fate decisions. Conceptually, slower-dissociating BCR interactions constrain GC recruitment/residence and limit needless SHM when narrow, primary responses suffice (“if it ain’t broke, don’t fix it”), whereas faster-dissociating GT immunogens like GT7 sustain GC fitness for pathogens requiring exceptional breadth like HIV-1 as well as other diverse pathogens like influenza, coronaviruses, and HCV.

Consistent with the intrinsic “traffic flow” model, GC fitness appears largely independent of extrinsic variables, a conclusion supported by multiple *ex vivo* and *in vivo* readouts. Short-term transcriptional profiling of CH31 UCA^hom/homdKI^ B-cells stimulated with eOD-GT7 versus eOD-GT8 np’s revealed comparable proliferative clusters but selectively enhanced GC programming with GT7 (**Fig 2**) in splenocyte suspension cultures lacking exogenous mitogens and organized follicular T–B zones under CD4-dependent, rate-limiting conditions. Importantly, these differences reflected uniform per-cell transcriptional shifts rather than expansion of discrete subsets. *In vivo*, clonal expansion of CH31 UCA^+^ B cells at peak GC occupancy was similar between eOD-GT7– and eOD-GT8–primed animals, even at both physiological and ultra-low precursor frequencies—conditions imposing limiting BCR availability rather than antigen or T cell help (**Fig S12**)—yet GC magnitude consistently favored GT7. DZ/LZ ratios and kinetics were also unchanged (**Fig S13**), indicating that intrinsic BCR signaling kinetics govern GC recruitment efficiency while preserving uniform DZ-LZ cycling, consistent with transcriptional programs observed *ex vivo*. Together, these findings identify GC fitness as a clonotype-specific intrinsic property determined by paratopic sensing of eOD-GT7 versus eOD-GT8 interactions and suggest that key B-cell fate programs-including GC competence-are largely dictated by intrinsic BCR properties despite extrinsic selection-modulating cues. This framework supports GT strategies that explicitly account for clonotypic requirements of tractable bnAb class members and suggests that culture-based systems could accelerate classification of bnAb clonotypes across Env epitope classes.

The divergence between proximal BCR activation readouts and those defining optimal GC priming *in vivo* underscores the need for culture systems that capture *bona fide* GC differentiation. Intrinsic clonotypic sensing of GT7 versus GT8 np’s—reflected by comparable proliferation but selectively enhanced GT7-driven GC programming—is concordantly observed during *in vivo* priming and within our *ex vivo* CH31 UCA^hom/hom^ ^dKI^ culture platform (**Figs 2,4**), supporting the latter’s use as a predictive and scalable modality for GT immunogen evaluation. Although KI models expressing bnAb precursors have transformed preclinical testing by enabling controlled analysis of precursor activation, clonal expansion, and GC recruitment^53–55,62,65^, iterative *in vivo* testing even in this setting remains resource intensive. Moreover, heterogeneity in precursor V(D)J clonotypic baselines across bnAb classes indicates that GT vaccination must inherently be precision-guided and thus requires predictive biophysical parameters linking antigen-specific design to GC fitness. Without high-throughput screening systems, immunogen optimization remains time/cost-prohibitive. The platform described here enables rapid ranking and down-selection of candidates using defined BCR metrics and go/no-go criteria (**Fig 6B**), prioritizing immunogens most likely to recruit lineage-specific precursors into GCs and promote improbable SHM events while minimizing *in vivo* validation and progressively accelerating clinical translation. As a prototype, the system could be further sensitized by incorporating hyper-VDJ mutagenic backgrounds like Rev3_L->F_ KI mice expressing a hyperactive polymerase-ζ variant^97^, which in CH31 UCA×Rev3_L->F_ compound KI models further accelerates SHM and indel accumulation at the hypermutable V_H_1-2-bearing CH31 clonotype across all base pairs, both post-prime and in full regimens^98^, increasing detection of breadth trajectories such as HC insertions.

In addition to accumulation of rare point mutations relevant for CH31 breadth, we previously demonstrated that multi-residue insertions in HCDR1 and HFRW3 required for Env flexibility/breadth^17,18^ can be reproducibly elicited in CH31 V_H_1-2 KI models through stepwise GT trimer immunization^19^. Here, we extend these observations by capturing a broad spectrum of indels during optimal eOD-GT7 np–dependent GC priming at physiological precursor frequencies (∼1/10^5^), with particular enrichment in HCDR1 and HFRW3 deletions. Intriguingly, these products have a remarkable resemblance to inferred first order predecessors of insertions identified in evolutionary reconstructions of the CH30–CH34 patient lineage^18^ and vaccine-induced clades in CH31 UCA dKI mice^19^, suggesting a near-mature breadth trajectory may be possible in as few as two immunization steps. Furthermore, relative to GT8 priming, GT7 also induced greater SHM accumulation at AID hotspot clusters flanking HCDR1 and HFRW3 regions, also patterns previously implicated in insertion acquisition during natural infection^18^ and stepwise GT vaccination^19^, and further supporting these events as *bona fide* selected (in-frame) pre-indels rather than stochastic byproducts. Thus, translationally, the estimated abundance of VRC01/CH31-class V_H_1-2 precursors estimated in humans (∼10^5^-10^6^)^43,64^ together with measurable frequencies (∼0.8%) of eOD-GT7-induced CH31 UCA^+^ B-cells in GCs harboring pre-insertion events **(Fig 5H)** suggests a potential accelerated pathway to CD4bs breadth that may partially substitute for prolonged accumulation of point mutations, particularly under strong recall selection. Convergent evidence from early clinical evaluation of a related CD4bs GT candidate^99^ and from HCDR3-dominated CD4bs regimens inducing breadth-associated HCDR2 insertions^100^ indicates that indel-enabled maturation may represent a generalizable feature of multiple CD4bs lineages for immunization to exploit. More broadly, these observations support the concept that vaccine immunogens can be rationally tuned not only to select specific precursors but also to bias evolutionary mutation pathways during GC maturation. They further suggest that CH31 precursor targeting could be incorporated into combinatorial regimens with complementary precursors like that for CH01-04, a lineage from the same donor belonging to the V2 apex-directed bnAb class from the same donor with fewer SHM to breadth but faces a distinct priming bottleneck related to clonotypic repertoire availability^39,40,101–104^. Such multi-lineage strategies may enable cumulative pathways to breadth. Ongoing CH31→CH01 transfer studies, together with developing analogous culture systems for additional GC fitness–amenable bnAb precursors/targets, should provide a framework for rationally designed GT prime–recall strategies capable of progressively assembling bnAb breadth across multiple Env epitope classes.

It will be fascinating to determine whether the priming parameters identified here for optimal GC activation and affinity maturation are similar in following immunization rounds, for example, during initial (shaping) boosting when recall likely will need to extend GC fitness, versus terminal boosting, when the GC response needs to be turned off. In this regard, memory phase constraints - including distinct signaling properties of IgG versus IgM/IgD BCRs - will require dedicated evaluation during shaping (SHM continuation) and polishing (response termination) phases^105^. Thus, while shared biophysical requirements for GC continuation, including minimal association rates^60,61^ and moderate affinity antigens^106^ likely extend from priming to boosting, additional parameters, such as platform heterogeneity in prime-boost regimens may further influence outcomes^107^.

Finally, regarding host controls across various HIV targets, this study reveals the biophysical rules that reliably trigger bnAb precursors defining BCR kinetic parameters crucial for their recruitment to - and retention in - GC reactions, focusing on the authentic CH31-CH34 lineage UCA, a prototypical CD4 mimic clonotype which is a more tractable HIV vaccine target in having human-relevant precursor frequencies^89^, capacity for breadth-conferring HC insertions^19^, and similarity to GC-competitive, only mildly polyspecific peripheral B-cells^23,24,71,72^ in humans^11,108^ But what about for bnAb precursors to other HIV targets? Notably, recent priming studies we have conducted using a completely distinct KI->WT transfer system to study the more signaling-responsive V2 apex targeting CH01 lineage UCA – recapitulates the need for higher antigenic valency combined with moderate affinity for optimal CH01-specific GC responses and serum cross-neutralization^101^. Furthermore, since most KI models expressing bnAb precursor V(D)J clonotypes other than the CH31/VRC01 ie CD4 mimic class have reduced (not increased) baseline BCR densities^62,63^ with accompanying varied, increased degrees of anergy-mediated GC exclusion^3,29,63,92,109^ and signaling incompetence^37,38,66,69,76,110^, it is unsurprising that they too, would have high valency requirements, also consistent with evidence that GC LZ B cells downregulate BCR densities to enforce stringent affinity discrimination via multivalent antigen presentation, independent of clonotype-specific thresholds^111–114^. By contrast, baseline signaling heterogeneity in these more anergic bnAb precursors would be expected to more significantly impact minimal k_a_ threshold, given increased valency alone only partially rescues GC responses in strongly anergic precursors^76,37,66,69^, and thus suggesting such lineages may additionally require higher association rates, or in some cases, may be “irredeemable”^23,24^. Encouragingly however, ‘HCDR3-binder’ GT MELs capable of eliciting cross-neutralizing CD4bs-specific serum responses^100^, demonstrate that appropriate kinetic tuning - including engineering for sufficient k_a_ - can overcome such constraints, even in more profoundly anergic lineages like CH103-CH106 ^31,38^. These findings thus support a broader model in which optimal bnAb-inducing strategies will require coordinated GT across several Env epitopes to maximize both bnAb precursor coverage and serum breadth. Given this clonotypic heterogeneity, future work should thus empirically tune clonotype-specific signaling range using the integrated *ex vivo* and *in vivo* prime testing systems in **Fig 6B** to eventually confirm the predictive framework outlined in **Fig 6C**. This proposed framework is also currently being further confirmed by incorporating CD4bs and V2 apex GTs into available protocols, to verify impact on serum neutralization breadth and on long-lived memory IgG^+^ B-cell maturation after terminal recall. In this regard, we have recently used the more anergic CH235 UCA dKI system (relative to the CH31 UCA dKI system)^19^ to demonstrate that comparably lower affinity, faster dissociating immunogens most potently drive C4bs-specific serum cross-neutralization and key rare SHM events in CH235 precursors^115^.

## Supporting information

Main text and figures

## ACKNOWLEDGEMENTS

We thank Dr. Vaughn Smider, CEO at ABS and Dr. Barton Haynes, Director at DHVI for providing facility resources at respective sites and for scientific advice. We thank Bill Schief and Chris Cotrell at Scripps for providing eODGT proteins. We also thank the DHVI protein expression facility (Director: Kevin Saunders), the LJI Microscopy and Histology facility (RRID:SCR_014835) for expert sample preparation, the LJI and UCSD NGS sequencing facilities, and the ABS/DHVI finance and administrative teams, for their support. Research reported in this submission was supported by the National Institute of Allergy and Infectious Diseases (NIAID) of the National Institutes of Health (NIH) under Award Numbers R01 AI087202 (LV) and R01 AI145656 (SMA and LV). The content is solely the responsibility of the authors and does not necessarily represent the official views of the NIH.

## AUTHOR CONTRIBUTIONS

K.A. performed SPR binding assays and immunogen QC. J.Z. and M.O. performed flow cytometry. A.G. performed immunizations and collected samples. A.G. and D.R. provided animal care J.Z. carried out B cell sorting, culturing, and *ex vivo* cellular analyses. J.S. generated and validated 10X single cell WGT and Ig-seq splenic libraries. Z.M. supervised immunofluorescence experiments and analyzed microscopy data. J.Z., M.O., and T.O. performed next generation sequencing. A.Z., T.O., M.D. and J.Z. provided bioinformatic analysis. M.D. and L.V. supervised research. S.M.A and L.V. provided resources and funding. J.Z., S.M.A, M.D. and L.V. conceptualized experiments and prepared the manuscript with input from all authors.

## DECLARATION OF INTERESTS

Provisional patents (LV and MD) are pending for the primary CH31 culture antigen testing platform, including the maiden version reported in this study.

## METHODS

### Antigens

eODGT6 and eODGT8 monomers were produced at the Duke Human Vaccine Institute, while eODGT7 monomers were produced at Scripps Research, all as previously described ^33^. eODGT6 60mer, eODGT7 60mer, and eODGT8 60mer np’s were also produced at Scripps Research based on published methodologies ^33^. Biotinylation and subsequent tetramerization of eODGT monomers has also been recently described ^61^. Briefly, Env sequences were first expressed with a C-terminal avidin tag (AviTag: GLNDIFEAQKIEWHE) using a BirA biotin-protein ligation kit (Avidity), after which excess biotin was removed using 10kDa MW spin columns (Amicon). Tetramerization was accomplished with stepwise addition of streptavidin (Invitrogen), using a 4:1 molar ratio of protein to streptavidin, to maximize for streptavidin site occupancy.

### Surface plasmon resonance (SPR)

SPR details have been extensively detailed previously ^61^. Briefly, SPR derived kinetic rates (ka and kd) and affinity measurements (KD) of monomeric eODGT6, eODGT7, and eODGT8 against the CH31 UCA IgG mAb were obtained using the Biacore S200 (Cytiva) instrument. A Protein A chip (Cytiva) was used to capture CH31 UCA IgG mAbs to a level of 200-300RU on flow cells 2, 3, and 4 for all proteins, while a negative control antibody, CH65, was captured onto flow cell 1 to approximately 300- 400RU for reference subtraction. Proteins were injected over the sensor chip surface using sequential injections of samples at various concentrations, followed by a 600s dissociation and regeneration with a 20s pulse of Glycine pH1.5. The Biacore S200 Evaluation Software (Cytiva) was used to analyze results. Binding to CH65, as well as buffer binding were used for double reference subtraction and accounted for both non-specific binding and signal drift. Curve fitting analyses were performed using the 1:1 Langmuir model.

### Mice

A colony of CH31 UCA^hom/hom^ ^dKI^ (V_H_DJ_H_^+/+^ x VκJκ^+/+^) mice ie, fully homozygous gl-CH31 KI mice on the C57BL/6j CD45.2 background were maintained and genotypically verified for all experiments in this study and have been previously described ^19^. B6.CD45.1 mice (used as recipients in adoptive transfer studies described further below) were purchased from The Jackson Laboratory (B6.SJL-Ptprca Pepcb/BoyJ, Strain #:002014). All animals reported in this manuscript (ie, either for *ex vivo* experiments or to generate reconstitution cohorts for vaccine priming studies) were 8-12 weeks of age, with equal numbers of males and females distributed across all experimental groups. All strains were housed in the ABS vivarium under a pathogen-free environments, at 12h light/dark cycles and at 20–25°C, in accordance with NIH guidelines. All animal procedures performed were approved by ABS Institutional Animal Care and Use Committee (IACUC)-approved animal protocols under AAALAC guidelines.

### Adoptive transfers and immunizations

After obtaining single-cell suspensions from CH31 UCA^homhom^ ^dKI^ spleens by mechanical dissociation, red blood cells were removed by ACK lysis, and B cells were purified by negative selection using the StemCell EasySep™ Mouse B Cell Isolation Kit, Cat #19854) according to the manufacturer’s instructions. Both viability and purity of B cells was routinely above 95%, as determined by live/dead and combined B220 & CD19 staining, respectively, while >85-90% of purified B-cells consistently carried both the knocked in HC and LC CH31 UCA V(D)J rearrangements, as verified by staining using a labeled eODGT8-60mer probe. CH31→WT chimeras were generated by retroorbital transfer of various amounts of purified donor (CD45.2+) CH31 UCA dKI B cells (numbers varying, depending on specific experiment) into recipient B6.CD45.1 mice, based on empirically determined input→output ratios (described in further detail in the results section, as part of validating this transfer model).

Primes with various eODGT proteins were performed 24 hours after transfer. 18 μg of each immunogen was diluted in DPBS and mixed with 60 μg of poly I:C (InvivoGen, Cat #tlrl-pic) in 200 μL total volume per animal. A minimum of 5 mice per group had this mix administered intra-peritoneally as a single-site injection using a 25G needle. For our initial immunization study specifically (which examined the impact of precursor frequency on eODGT immunogen abilities to induce GC responses and SHM), we also reconstituted additional mice at each frequency (ie, extras, relative to those used for immunizations), to verify output frequencies of CH31 UCA dKI+ B-cells in recipient spleens 24h post-transfer, by staining with both CD45.2 and the eODGT8-60mer bait.

### Proximal BCR signaling analysis in primary B-cells (Ca^+^ flux & phospho-kinase flow assays)

For analysis of the elevation of [Ca^2+^] cytosol induced by transient BCR-signaling, RBC lysed CH31 UCA dKI and (where applicable, as controls) WT/B6 splenocytes were stained with BV650 anti-B220 and R78 anti-CD19 (Cat#567063) for 40 minutes. After washing with HBSS, pre-stained cells were loaded with Fluo-4 via by mixing with an equal volume of Fluo-4 loading solution (Fluo-4 Direct Calcium Assay Kit, ThermoFisher, Cat# F10471). After 30 min incubation at 37 °C and then again at RT, cells were washed with HBSS and incubated with LIVE/DEAD Near-IR for 30 minutes. Cells were washed twice with HBSS, resuspended in [Ca^2+^]-supplemented HBSS, incubated at room temperature for 5 minutes, then transiently activated by various eODGT immunogens or F(ab’)_2_ anti-IgM (Southern Biotech). Raw Fluo-4 MFI data was acquired on a BD Aria III flow cytometer and total B-cell (B220+CD19+) responses were analyzed by FlowJo software. Calcium flux results were then quantified using Microsoft Excel and GraphPad Prism v9, as previously described ^61^. Briefly, after subtracting for autofluorescence, Fluo-4 MFI was normalized with respect to the maximum signal of the IgM control and calcium flux values were presented as a percentage (% of F(ab’)_2_ Anti-IgM Max).

For phospho-flow kinase analysis of primary B-cells, RBC-lysed splenocytes from 8-12 week CH31 UCA dKI (or, as controls) WT/B6 (strain-matched) mice were subjected to negative B-cell separation using the StemCell EasySep™ Mouse B Cell Isolation Kit. Purified B-cells were stained with Live/Dead NIR in DPBS, followed by surface staining with BV650 anti-B220 and BB515 anti-CD19 (BD Cat#564509) in BD Brilliant Stain Buffer. Cells were then washed and stimulated in HBSS (ThermoFisher, Cat#14025134) with eODGT immunogens for different times as indicated in the results. Reactions were then stopped by immediate addition of BD Cytofix™ Fixation Buffer (BD Cat#554655), vortexed, incubated for 10-15 minutes at 37°C, spun down at 1000g for 5min. After supernatant was removed, cells were then permeabilized (BD Phosflow™ Perm Buffer III, Cat. No. 558050) on ice for 30 minutes, washed with Stain Buffer (FBS), stained with Alexa Fluor® 647 Mouse anti-Btk (Cat. No. 558528) and PEcy7 anti-Syk (Y352) (Cat# 561458), and protected from light, overnight at 4°C. Data were acquired using a BD Aria III flow cytometer and analyzed using FlowJo software (v.10.8.1). Normalized MFI for each timepoint was calculated as the ratio of treated sample MFI relative to unstimulated sample MFIs.

### Flow-cytometric B-cell phenotypic analysis

For quantifying GC B-cell subsets in eODGT-primed CH31^UCAhom/homdKI^→WT chimeric animals, spleens or lymph nodes were sliced into small pieces, and passed through cell strainers. RBC lysed cells were stained with Live/Dead NIR for 30 min, then further stained with BD Brilliant Stain Buffer containing eODGT8-PECF594-tetramers and fluorochrome-labeled mAbs. Primary labeled mAbs (all from BD Biosciences) were used at 0.5 μg/ml and included BV421 GL7 (Cat# 562967), BV510 anti-IgG1(Cat#740121), BV510 Anti-IgG2a/2b (Cat#744293), BV510 Anti-IgG3 (Cat#744134), BV605 anti-CD45.2 (Cat#563051), BV650 anti-B220 (Cat^#^ 563893, clone RA3-6B2), FITC anti-CD45.1 (Cat#553775), BB700 anti-CD138(Cat# 742124), PEcy7 anti-IgM (Cat# 552867), APC anti-IgD (Cat# 560868), and R718 anti-CD38 (Cat#752576).

For detecting distal BCR signaling outcomes (ie early proliferative, activation and GC differentiation signatures) in short-term CH31^UCAhom/homdKI^ splenic cultures, whole RBC-lysed splenocytes from CH31 UCA dKI mice were pated in 96 well plates at 5×10^6^/ml for 24h at 37°C 5% CO_2_ in complete RPMI 1640 medium (ie, supplemented with 10% FCS, 100 μM 2-Mercaptoethanol and 50 U/ml penicillin/streptomycin), either in media alone (ie, with no ODGT immunogen stimulation), or in the presence of eODGTs, at indicated concentrations and valencies. B-cells were stained with Live/Dead NIR in DPBS, followed by surface staining for capturing all B-cells using BV650 anti-B220 and BV786 anti-CD19 (Cat#563326), for assessing their early-intermediate activation/proliferative status using BV421 anti-CD25 (Cat#562442) or PEcy7 anti-CD69 (Cat#561928), and for evaluating their GC differentiation status using PE anti-GL7 (Cat#555361) and BV510 anti-mouse MHC class II I-A/I-D alloantigenic determinants (Cat#742893).

### CH31 UCA^hom/hom^ ^dKI^ culture single cell genome-wide transcriptomic libraries and analyses

RBC lysed splenocytes (>20 million per treatment group) from up to three pooled CH31^UCA^ ^hom/hom dKI^ mice were plated in 24 well plates at 5×10^6^/ml for 24h in complete RPMI 1640 medium alone or in the presence of 1 nM eODGT7 or eODGT8 np’s or 20ug/ml F(ab)_2_ anti-μ. Cells were harvested, washed with 1xDPBS and single cell genome-wide transcriptomics libraries were constructed using the 10X Chromium Single Cell 5’ v2 workflow protocol (CG000330, Rev F)and a Chromium Next GEM Single Cell 5’ Gel Bead Kit v2 (16 rxns PN-1000263). Briefly, master mix, Chromium Partitioning Oil, and a Chromium Next GEM Chip K was used to partition thousands of single cells into a pool of Gel of Beads-in-emulsion (GEMs), containing approximately 750,000 10X barcodes, where each cell was labeled with a unique barcode to index its transcriptome. Cells were saturated with 10x barcoded gel beads so that >90% of generated GEMs lacked cells to ensure single resolution per 10x barcoded gel bead. 10X-barcoded single splenocyte-containing GEMs were then loaded onto a 10x Chromium X controller to carry out partitioning by emulsification at a nanoscale level. 10x barcoded cDNA was converted from poly-adenylated mRNA by dissolving co-partitioned 10x barcoded gel beads and cell lysates, pooled and purified from excess 10x reagents and non-cDNA cell debris with Chromium recovery agents and silane magnetic MyOne (Dynabeads prior to cDNA amplification. Amplified cDNA was subject to a one-sided size selection SPRI bead cleanup (SPRI^select^ Reagent Kit, Beckman Coulter), verified for size distribution using an Agilent 2100 Bioanalyzer, and quantified using a Qubit 4 Fluorometer. Transcriptomics libraries were generated using the 10x Library Construction Kit (16 rxns PN-1000215), in which 50ng of starting sample was fragmented, ligated to sample index adaptors, and subjected to index PCR (Dual Index Kit TT Set A), with sample indexes contained priming sites used by Illumina sequencers during sequencing, and further SPRI size selections carried out prior to adaptor ligation, and before and after sample index PCR amplification. Samples were sequenced in both directions on an Illumina NovaSeq 6000 (LJI NGS Core) with a cutoff of 20,000 reads per cell for each constructed sample transcriptomics cDNA library.

Transcriptomics information was obtained by generating C-Loupe files, using Cell Ranger hosted by the 10x Genomics cloud interface. FASTQ files were uploaded from 10x Genomics library sequencing and data was analyzed using the Cell Ranger pipeline (https://www.10xgenomics.com/support/software/loupe-browser/latest). Loupe Browser (Volume 8.0) was used for differential gene expression comparisons among splenic cells treated with α-IgM BCR F(ab’)_2_, eOD-GT7 np, or eOD-GT8 np. Untreated splenic cells (ie, cultured in medium alone) were used as controls for establishing baseline expression levels when needed. Main gene expression clusters identified for all four treatments were assigned to cell type based on expression of lymphocytic cell markers such as CD79 (α and μ) for B-cells, and either CD4 or CD8 co-expressed with CD3 for T-cells, and in Fig 2D, are encircled and annotated in the map to the left, with their patterns in specific treatment groups indicated to the right (B-cell and T cell gene clusters in Fig 2D are represented by green and gray shading, respectively). Gene expression clusters were initially compared against each other for differential gene expression, in order to establish B or T cell subsets as well as proliferation or activation status. FoxP3 was used to establish clusters enriched for regulatory T cells, as MKI67, Pclaf, and cell cycle gene levels were used to determine proliferation status of B cells. Differential gene expression among treatments or clusters was established using significant values of p<0.05 and/or repeated expression differences among different lymphocyte-specific gene clusters. Where baseline expression of a particular gene was significant as determined by gene levels in the untreated group, cutoff expression levels were set at or near the halfway point of the expression range to subtract baseline expression. Heatmaps were generated using all upregulated genes per treatment group or cluster in each comparison, followed by comparisons among genes of interest for a particular lymphocytic population. The list of genes of interest were generated *a priori* based on previous publications^116,117^; also see (https://www.bio-techne.com/research-areas/immunology/b-cell-development-and-differentiation-markers) and/or known genes relevant for specific immune functions (such as AICDA, CD19, etc..).

### Sorting of single donor-derived B-cells from GCs and Isolation of V(D)J pairs

Cell sorting was performed using a BD Aria III sorter (ABS flow facility). Briefly, CD45.1 depleted cells were collected using MagniSort™ Mouse CD45.1 Negative Selection Kit (Cat# 8802-6848-74) and stained. Donor GC B cells were visualized by first gating on lymphocyte singlets, followed by exclusion of doublets and LIVE/DEAD NIR stained cells (to discriminate live from dead cells), subsequent gating for B220^+^CD138^-^ (total B-cells), then CD38^-^GL7^+^(GC B cells) and finally, gating for CD45.2^+^CD45.1^-^ (Donor GC B-cells). Donor GC cells were two-way bulk sorted into Eppendorf tubes containing HBSS with 5% FBS. To recover V(D)J rearrangements from individual donor-sorted (CD45.2+) GC (GL7+CD38-) B cells, pellets were spun down for 5 min at 500g to remove excess HBSS/FBS. Cells were gently resuspended and single cell paired IgH/L library construction was carried out following instructions provided by 10X Genomics (CG000330, Rev F), using a 10x Chromium X Series controller, 10x Genomics Chromium Single Cell Mouse BCR Amplification Kit (16 rxns PN-1000255), and a Chromium Controller Single Cell RNA Sequencing Analyzer (10X Genomics). V(D)J libraries were generated using the 10x Transcriptomics Library Construction Kit (16 rxns PN-1000190), in which 50ng of starting sample was fragmented, ligated to sample index adaptors, and subjected to index PCR (Dual Index Kit TT Set A, 96 rxns PN-1000215), with sample indexes contained priming sites used by Illumina sequencers during sequencing, and further SPRI size selections carried out prior to adaptor ligation, and before and after sample index PCR amplification. cDNA and V(D)J library quality and quantity were checked using Qubit 4 Fluorometer and Agilent 2100 Bioanalyzer. Samples were sequenced in both directions on an Illumina NovaSeq 6000 (LJI NGS Core) using a 150 bp configuration, with a cutoff of 5000 paired reads per cell for each constructed sample V(D)J cDNA library.

### Immunogenetic & SHM sequence analysis in bnAb precursor donor-derived GC B-cells

Sequence analysis of CH31 UCA V(D)J rearrangements was performed based on previously described methods ^19^. Briefly, full-length reads were assembled, deduplicated, and annotated with immunogenetic information using Cell Ranger (v7.2). Reads that were identified as non-functional (e.g., out-of-frame, missing invariant Ig gene amino acids, or the presence of stop codons) were excluded from analysis. Frequencies of individual mutations from IgH/L paired V(D)J rearrangements of individual donor GC B-cells in each immunization group (or where applicable, each individual mouse) were calculated after aligning unique reads to the translated CH31 UCA sequence, using in-house bioinformatics programs including Lasergene software.

### Immunofluorescence analysis of bnAb precursor-derived B-cells in GCs

The fraction of donor (CD45.2+) CH31 UCA^hom/hom^ ^dKI^ B-cells in CH31 splenic GCs was determined following a previously established protocol ^53^ with the following specific modifications. Spleens were embedded in OCT and flash-frozen using the vapor phase of liquid nitrogen. Blocks were stored at - 80C and were sectioned at 5 um, skipping 100um between each section. Three sections from each block were adhered to TOMO slides and airdryed for 1h in a tissue culture hood. Slides were fixed with a 1:1 mixture of methanol and acetone chilled to -20C for 10 minutes and washed 2x for 10 minutes in PBS. Slides were placed in Freequenza racks (https://dx.doi.org/10.17504/protocols.io.3byl4ko9jvo5/v2), blocked for 1 hour with 0.5% BSA, 0.1% Tween-20 in PBS, and a 1/100 dilution of TruStain FcX (anti-mouse CD16/32, Biolegend 101320). Slides were incubated with 1:100 dilutions of the following directly-conjugated antibodies o/n at 4C (GL7-PE, clone GL7, Biolegend 144607, TCRβ-AF488, clone H57-597, Biolegend 109216, CD45.2-AF647, clone 104, Biolegend 109817, and B220-AF750, clone RA3-6B2, Novus Bio FAB1217S-100UG), washed 5 times with PBS, incubated with 50 ug/ml Hoechst for 10 minutes, washed 5 times with PBS, and coverslipped with Prolong Gold.

Whole slide images were acquired with 40x 0.95NA objective using Zeiss AxioScan Z1 equipped with Colibri7 LED illuminator, individual band-pass filter sets, and Hamamatsu Orca Flash4.0 v2 camera. Image analysis was done in QuPath 0.5.1 based on protocols in ^118^. Briefly, a pixel classifier was trained to detect tissue regions and annotations were manually corrected to remove artifacts. GCs were annotated with a pixel classifier trained on the GL7 channel, and objects were created and manually checked for accuracy. StarDist cell detection was performed in GCs and an object classifier was trained to detect CD45.2 cells.

### Statistical Analyses/Methods

Statistics were performed by either GraphPad (Prism) or Excel (Microsoft) software to determine p values by Mann Whitney U tests or where applicable, one-way ANOVA tests. P values between <0.05 (minimally significant) and <0.0001 (maximally signifcant) were used to determine significance levels.

## Notes

### Competing Interest Statement

The authors have declared no competing interest.

