## Supplementary material for "Binding dynamics shape germinal center broadly neutralizing responses to HIV priming": Main text and figures

### SUPPLEMENTAL MATERIALS

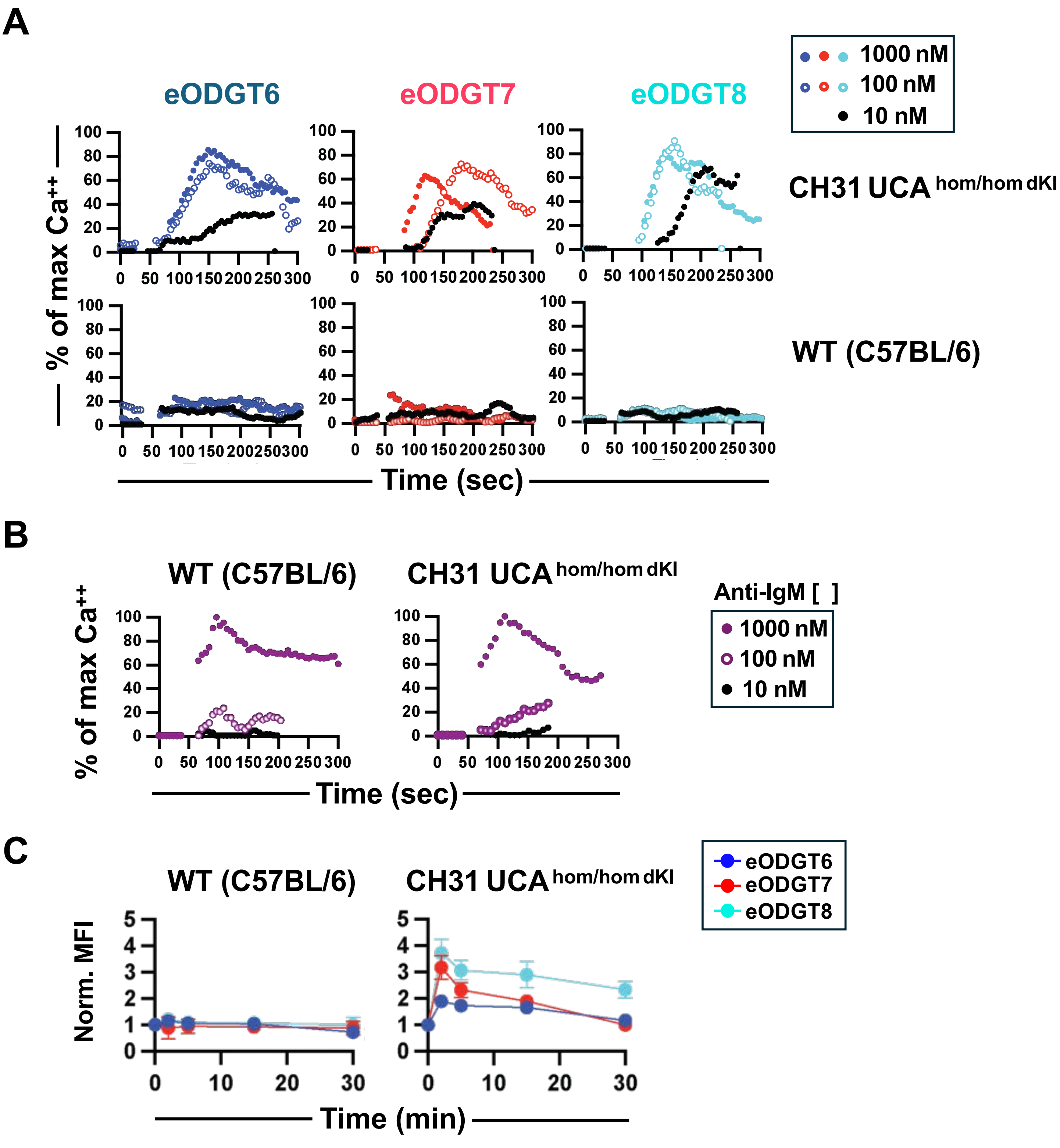


**Figure S1. Proximal *ex vivo* BCR signaling by CH31$^{UCAhom/homdKI}$ or C57BL/6 (WT) primary B-cells in response to eODGT nanoparticles. (A-B)** Graphical representation of flow cytometry-evaluated calcium flux kinetics in splenic B-cells directly harvested from naïve CH31$^{UCA\ hom/hom\ dKI}$ or wild type C57BL/6 (strain matched, non-KI control) mice. Primary splenic B-cells were stimulated with either **(A)** eODGT6, eODGT7, and eODGT8 60mer np immunogens or **(B)** F(ab')$_2$ anti-IgM at the varying concentrations denoted by inner legends. Results are presented as % of peak response to saturating concentration (1000 nM) of anti-IgM and is representative of two independent experiments with three mice/strain tested. **(C)** Graphs showing Syk (Tyr352) phosphorylation kinetics of CH31$^{UCA\ hom/hom\ dKI}$ or C57BL/6 primary splenic B-cells in response to eODGT 60mers. Data is represented as normalized MFI (ie fold increase over baseline MFI in unstimulated primary B-cells).

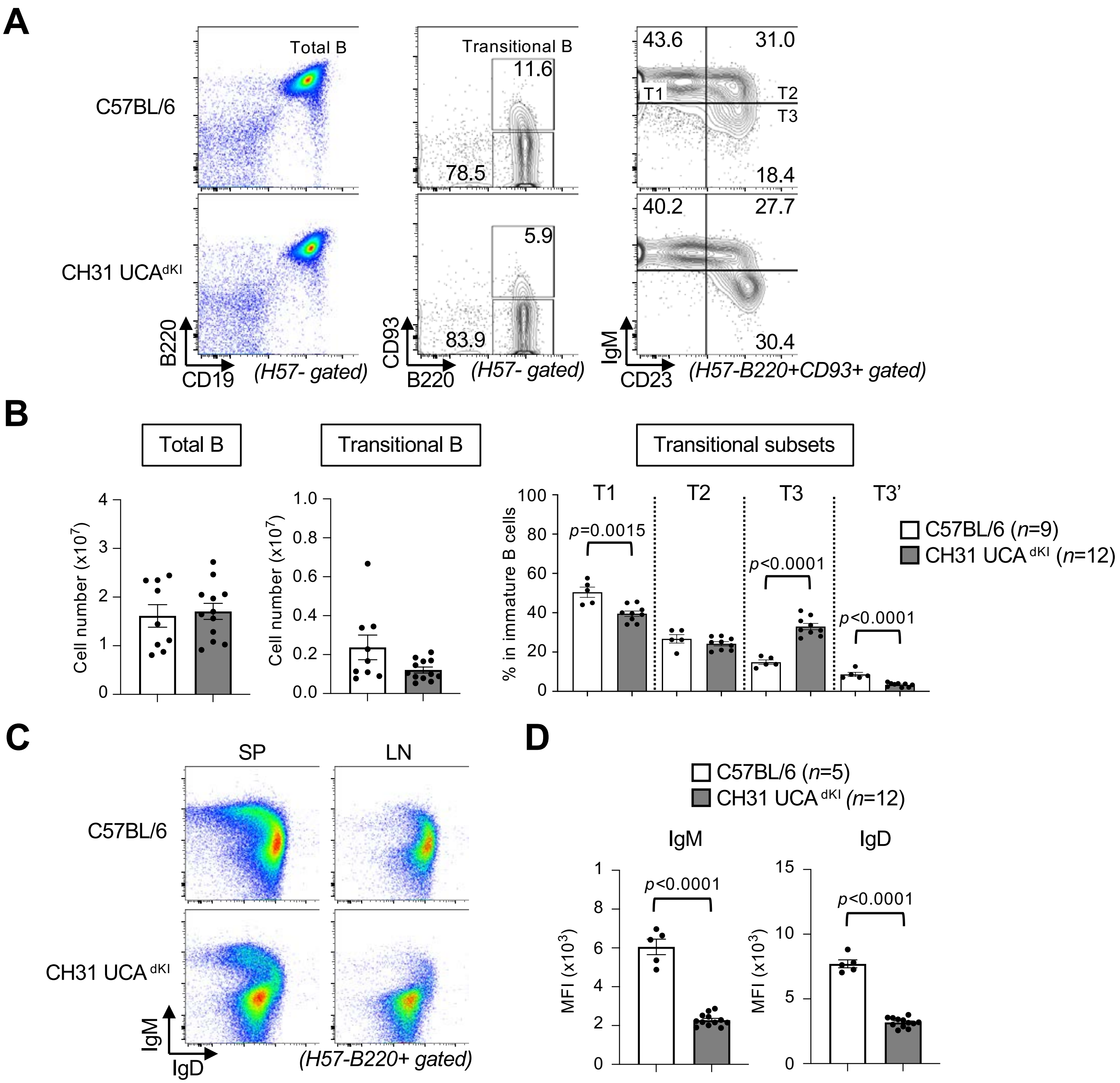


**Figure S2. CH31 UCA $^{dKI}$ mice have lowered BCR densities in peripheral lymphoid organs. (A-B)** Developmental analysis of B-cells in spleens of 8-12 wk naïve CH31 $^{UCAdKI}$ and strain-matched WT control (C57BL/6) mice. **(A)** Representative FACS histograms of splenic B-cell development, indicating percentages of live splenocytes that are total B-cells (CD3-B220+CD19+; *left* column) or Transitional B-cells (CD3-B220+CD93+; *middle* column). Transitional splenic B-cell subsets were further fractionated into T1, T2, T3 and T3' subsets, based on IgM and CD23 discrimination (*right* columns). **(B)** Graphical representation of flow cytometric determination of splenic B-cell percentages in populations shown in panel (A). Note the lack of significant differences between C57BL/6 and CH31 $^{UCA\,dKI}$ total and transitional B-cell populations, but the significantly expanded anergic (T3) splenic B-cell subset in CH31 $^{UCA\,dKI}$ mice. (**C**) Representative FACS histograms showing surface IgM and IgD BCR densities of B cells from spleen (SP) and lymph nodes (LN) of CH31 $^{UCAdKI}$ and C57BL/6 mice. **(D)** Graphical representation of surface BCR densities on total SP and LN B-cells, as analyzed in panel (A). Note the significantly downregulated surface IgM and of IgD BCR levels in total B-cells from CH31 $^{UCAdKI}$ mice. Data presented in panels (B) and (D) was graphed with GraphPad Prism 9 software and two-sided student's t-tests were used to perform parametric statistical comparisons.

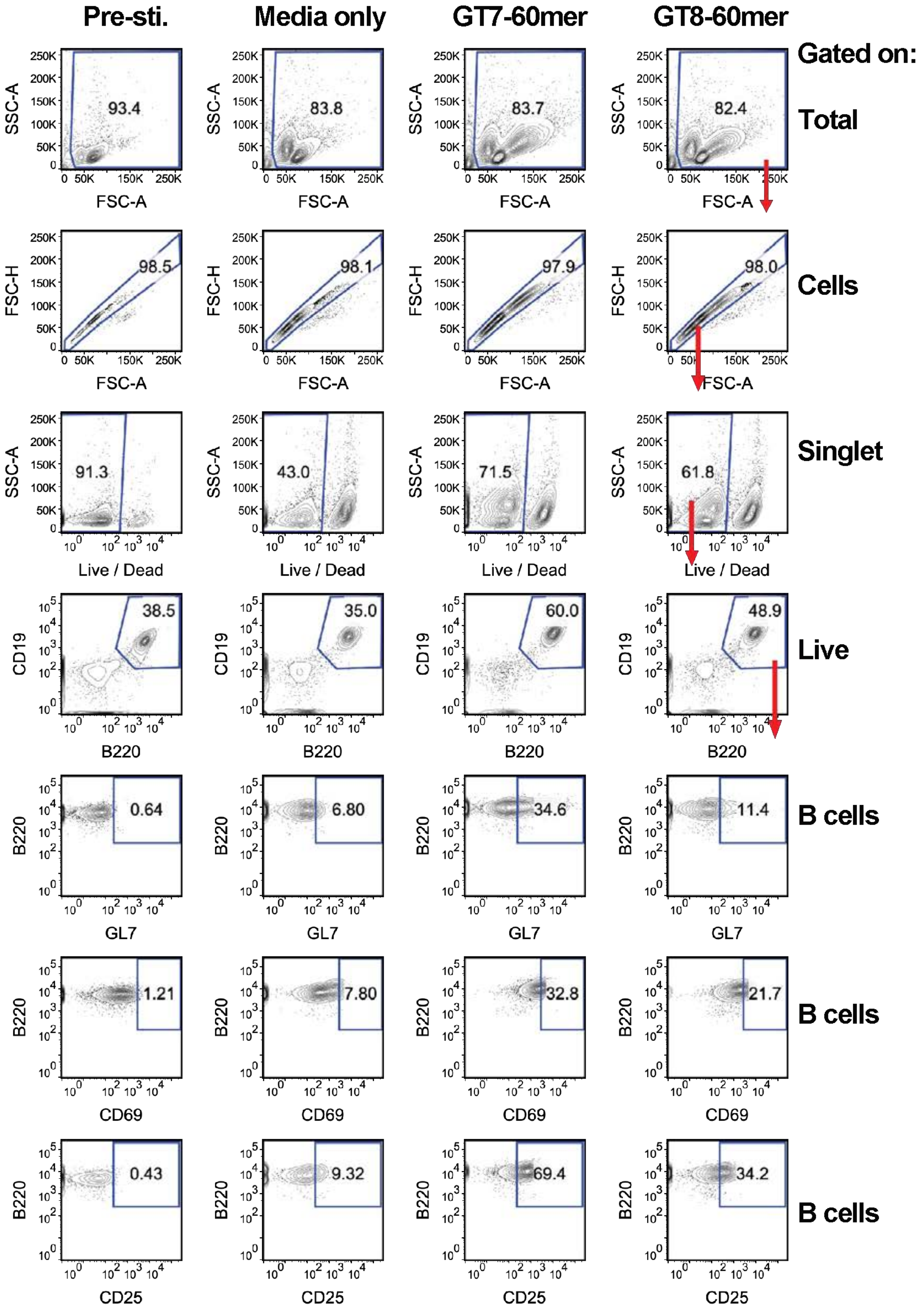


**Figure S3.** Flow cytometric gating scheme used to evaluate the percentage of total CH31 UCA $^{\text{hom/hom dKI splenic}}$ B-cells with upregulated surface expression of GL7, CD69 or CD25, after 24h in culture with 1nM eODGT7 or eODGT8 60mers. Shown are FACS histograms representative of a single culture wells for each stimulation group. The % of splenic B-cells with increased surface expression was calculated by first determining baseline levels for each marker ie, setting gates as close to 0% as possible at t=0 (prior to stimulation; left-most column) subtracted from percent of B-cells in these pre-set gates when cultured for 24h in media only (2$^{nd}$ column). Resulting baseline percentages were then used to subtract from those determined in pre-set gates in eODGT7 or eODGT8 np-treated groups.

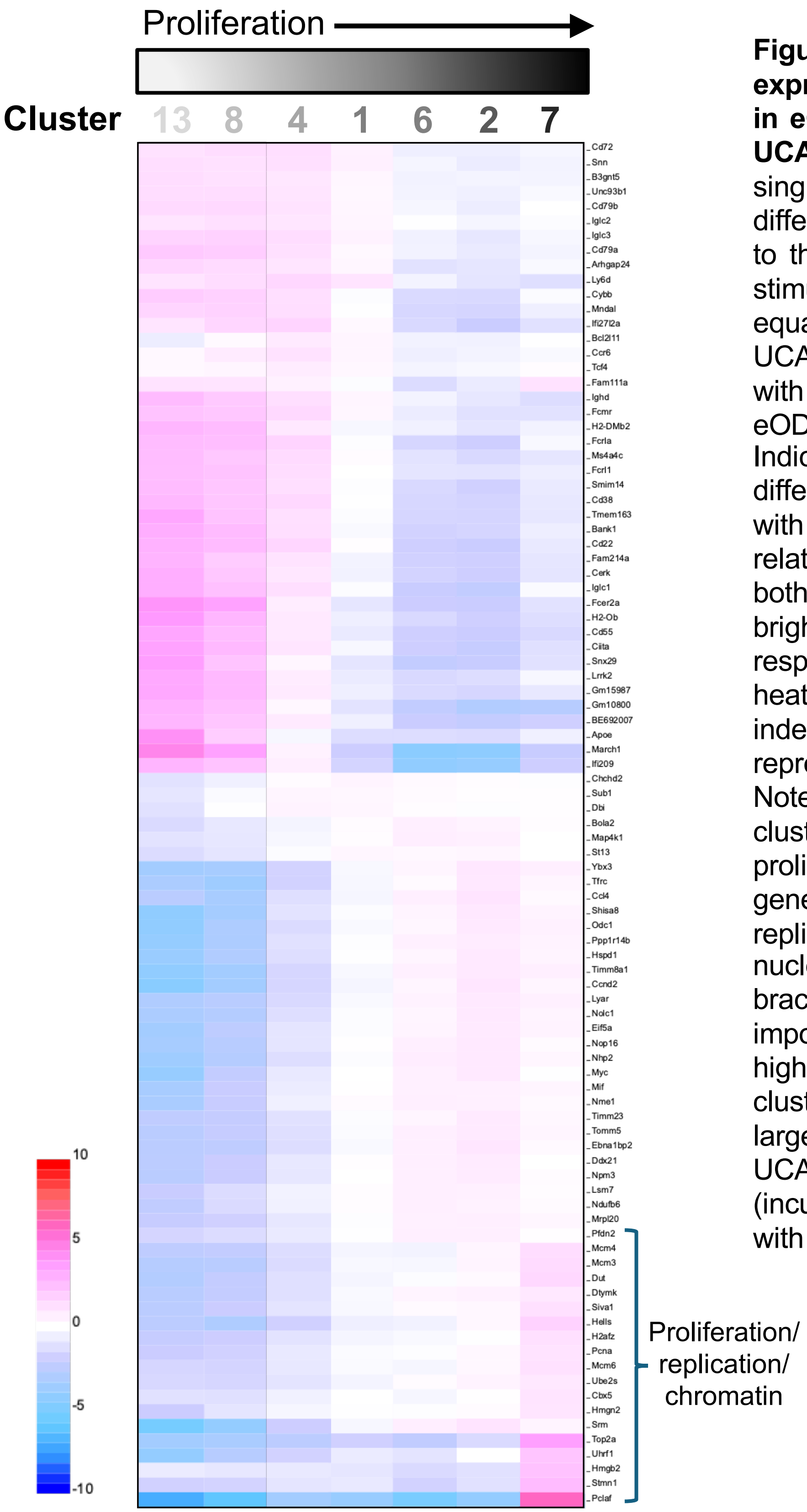


**Figure S4. Differential gene expression amongst B-cell clusters in eODGT np-treated splenic CH31 UCA$^{hom/hom\ dKI}$ cultures.** Shown are single cell transcriptome heat maps of differentially expressed genes (listed to the right) amongst eODGT7/8 np-stimulated B-cell clusters from an equal combination of CH31 UCA$^{hom/homdKI}$ splenic cultures treated with either 1nM eODGT7 np or eODGT8 np harvested for 24h). Indicated at the lower left is a differential gene expression index with fold increase or decrease in relative gene expression represented both numerically (-10 to +10) and by brightness of red or blue color, respectively. Also shown above the heat map is a B-cell proliferation index, with degree of proliferation represented by shading darkness. Note these analyses reveal that B cell clusters can be ordered by proliferation based on expression of genes involved in cell cycle, DNA replication, chromatin structure, and nucleotide metabolism (denoted by bracket at lower right position). Most importantly, the three clusters with the highest levels of these genes (ie, clusters 7, followed by 2 then 6), were largely absent from splenic CH31 UCA$^{hom/hom\ dKI}$ cultures left untreated (incubated in media alone) or treated with control anti-BCR reagent.

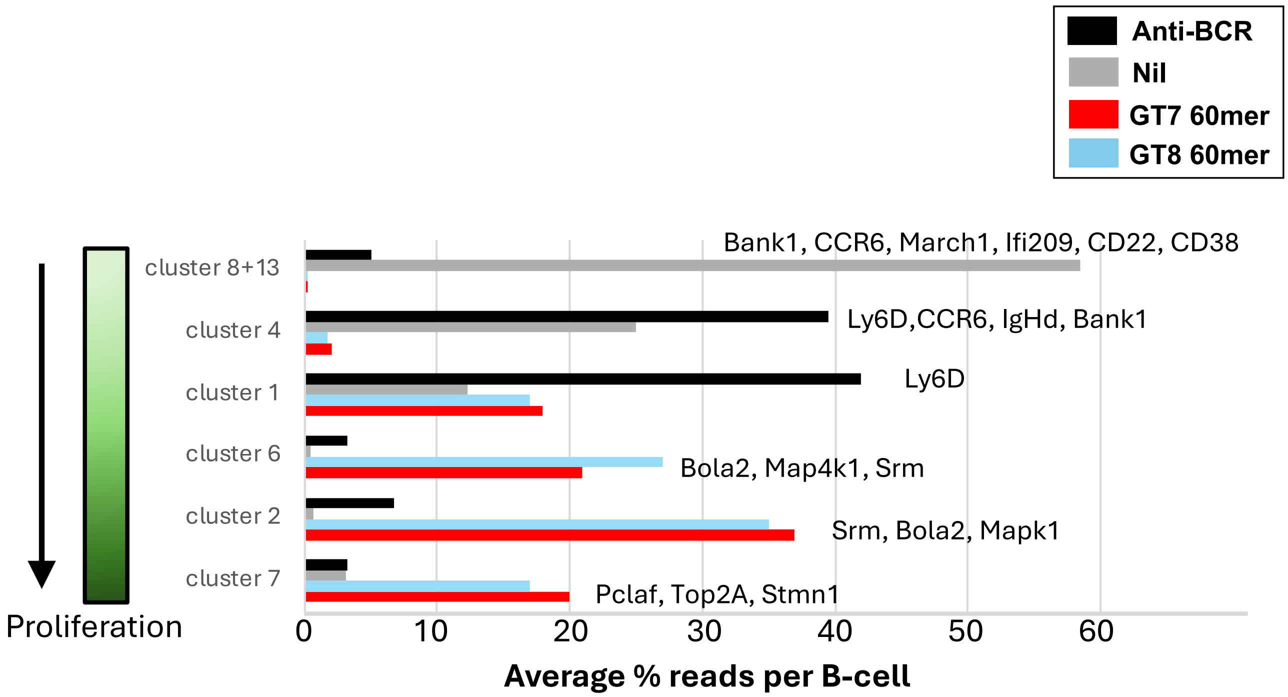


**Figure S5. Expression levels of overrepresented genes amongst B-cell clusters, shown by splenic CH31 UCA$^{hom/hom\ dKI}$ culture treatments.** Graphical schematic of preferentially expressed signature genes (listed to the right) amongst B-cell transcriptomic clusters from CH31 UCA$^{hom/homdKI}$ splenocytes cultured for 24h in media alone (nil; gray bars) or in the presence of either 1nM eODGT8 np, eODGT7 np, or anti-BCR (black, blue, or red bars, respectively). The x-axis represents the relative percentage of transcriptome reads per B cells expressing the denoted cluster-specific genes, for each treatment group. Note that eODGT7 np treatment induced the greatest fraction of gene-specific reads in B-cells found in cluster 7, ie putative early stage (DZ) GC B cells.

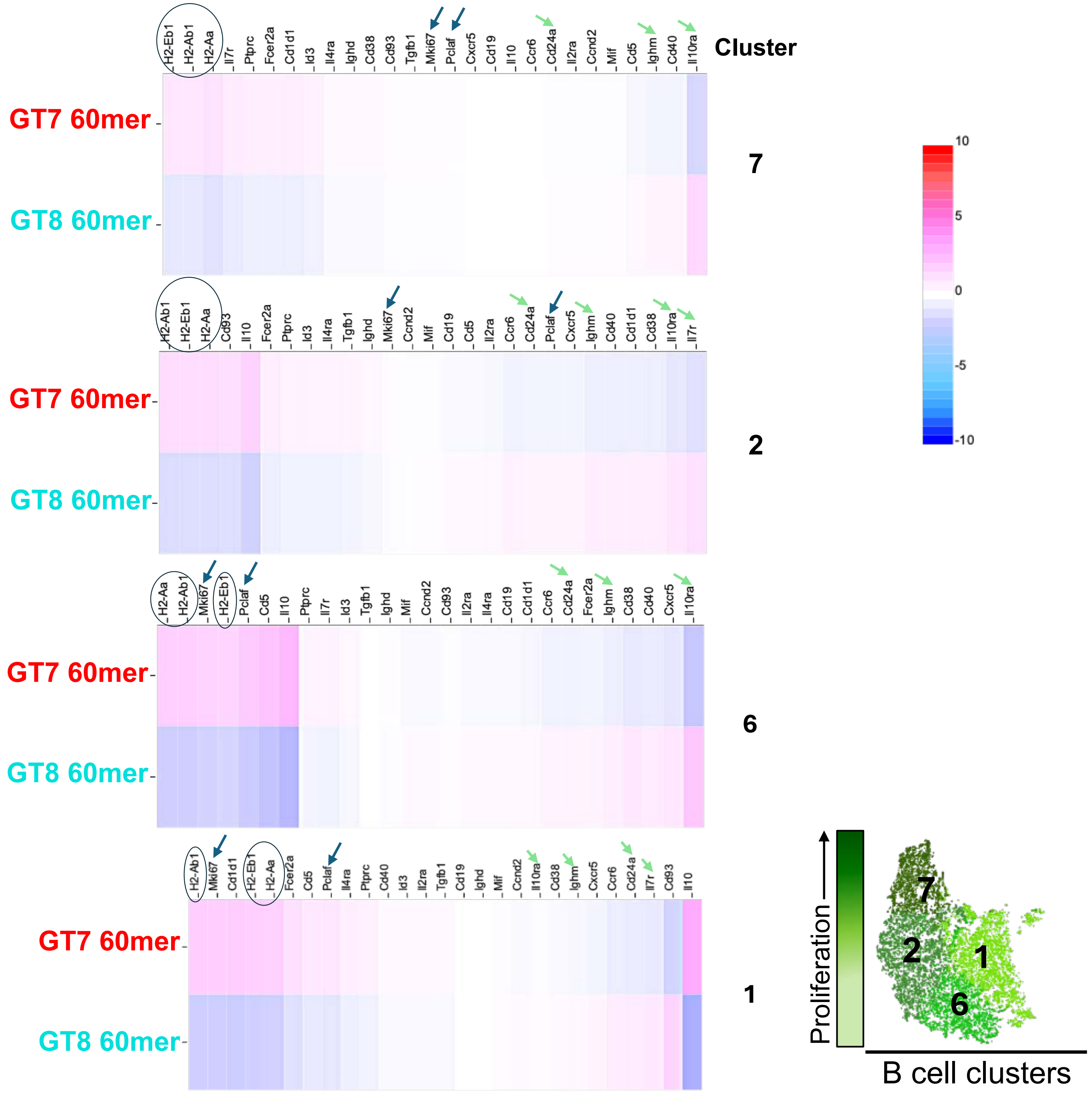


**Figure S6. Differential expression of select GC/Tfh-relevant genes between eODGT7 and eODGT8 np-treated splenic CH31 UCA$^{hom/hom\ dKI}$ cultures is conserved across proliferating B-cell clusters.** For each of the B-cell clusters amongst the more proliferative ones (1,2,6 and 7) are single cell transcriptome heat maps of genes differentially expressed between CH31 UCA$^{hom/hom\ dKI}$ splenocytes stimulated for 24h with 1 nM eODGT7 or eODGT8 60mers. Key GC-associated genes upregulated in eODGT7 np-treated cultures are highlighted, including MHC class II genes (circled) and GC-specific proliferation genes, (blue arrows), as well as genes upregulated in eODGT8 np-treated cultures like interleukin receptors associated with Th2/Tfh dampening (green arrows). Note that the differential expression patterns of many of these key genes is apparent even in clusters clusters 2,6 and 1, which exhibit progressively less proliferation/earlier differentiation relative to cluster 7. Indicated at top right is a differential gene expression index with fold gene up- or downregulation represented both numerically (-10 to +10) and by brightness of red or blue color, respectively. Also shown to the bottom right is a B-cell proliferation index (merge of eODGT7 & eODGT8 np-stimulated clusters) with cluster proliferation depicted by degree of green shading.

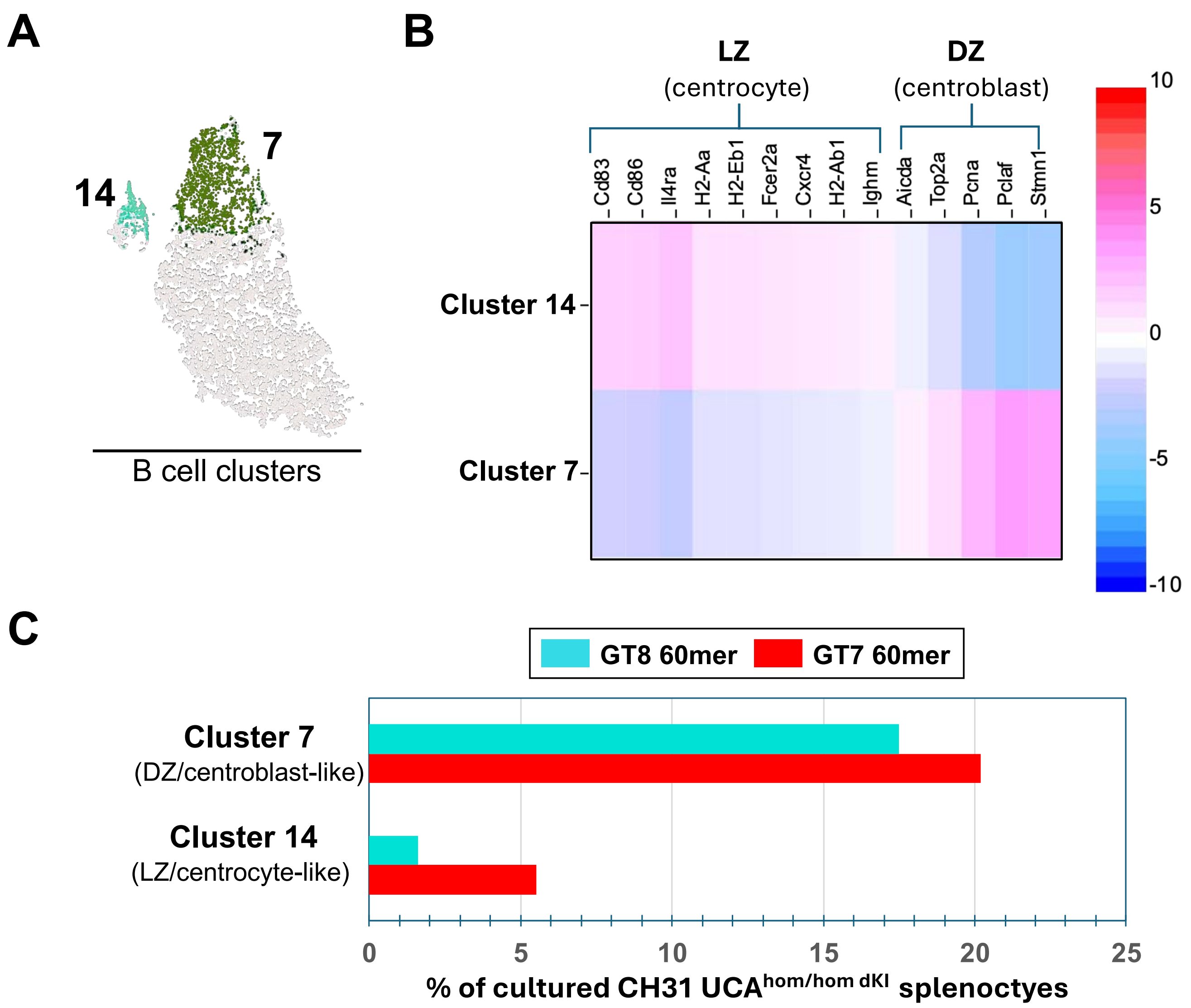


**Figure S7. GC Dark zone (DZ) centroblasts and Light Zone (LZ) centrocytes are likely represented by B-cell clusters 7 and 14, respectively. (A-B)** Combined eODGT7/8 np treated-culture transcriptome data was used to define putative novel (minor) B-cell cluster 14, in relation to (major) B-cell cluster 7. **(A)** Cluster map showing closest relative transcriptome similarity of GC LZ/centrocyte-like cluster 14 and DZ/centroblast-like cluster 7. **(B)** Heat map of signature a priori GC LZ and DZ genes differentially expressed between both GC-like clusters. Note that LZ/centrocyte-like cluster 14 has upregulated expression of genes involved in Tfh synergy like MHC class II, CXCR4, IL4 receptor, CD83, and CD86, whereas DZ/centroblast-like cluster 7 instead has upregulated expression of genes involved in SHM and proliferation such as Pclaf, PCNA, and AID. Indicated to at the right is a differential gene expression index with fold gene up- or downregulation represented both numerically (-10 to +10) and by brightness of red or blue color, respectively. **(C)** Fraction of total eODGT7 np-stimulated splenic CH31 UCA$^{hom/hom\ dKI}$ cultures whose genes originate from DZ-like cluster 7 and LZ-like cluster 14. Note the especially expanded fraction of B-cells originating from eODGT7 np-treated cultures relative to their eODGT8 np-stimulated counterparts.

**A**

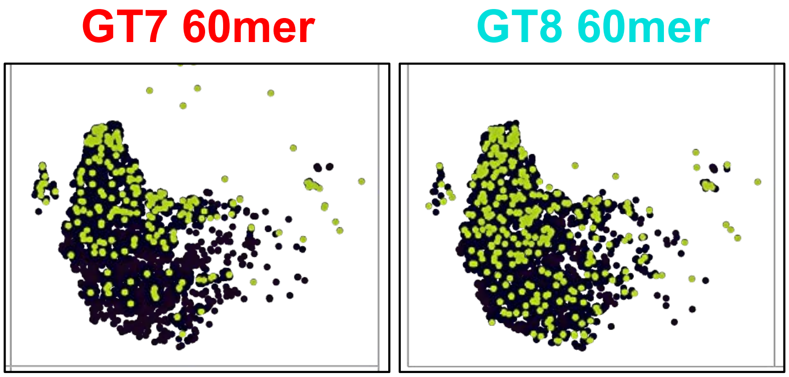


**B**

| Gene Name | Fold increase/ B10-like cell | P-value |
|---|---|---|
| IL10Rα | 12.73 | $3.95 \times 10^{-7}$ |
| SLC8A1 | 3.25 | $4.79 \times 10^{-4}$ |
| Gm13546 | 2.83 | $1.81 \times 10^{-3}$ |
| RAMP3 | 3.32 | $1.95 \times 10^{-3}$ |
| TBC1D8 | 2.79 | $2.73 \times 10^{-3}$ |
| CCL5 | 2.93 | $2.73 \mathrm{X} 10^{-3}$ |
| CACNB3 | 3.18 | $3.54 \times 10^{-3}$ |
| ADAM23 | 3.63 | $3.76 \times 10^{-3}$ |
| CXCL16 | 2.65 | $5.91 \times 10^{-3}$ |
| FSCN1 | 3.09 | $1.33 \times 10^{-2}$ |
| TBC1D4 | 2.48 | $1.33 \times 10^{-2}$ |
| CCR2 | 3.39 | $1.61 \times 10^{-2}$ |

**Figure S8. Expansion of a distinct $B_{Reg}$/B10-like subset of B-cells with an upregulated Th2/Tfh-dampening program in eODGT8 np-treated CH31 UCA$^{hom/hom\ dKI}$ splenic cultures. (A)** Single cell gene expression maps showing (in green shading) profiles of IL-10Rrα (associated with $B_{Reg}$ Tfh2-repressing/Th1 responses) across all B cell clusters in CH31 UCA$^{hom/hom\ dKI}$ splenic cultures stimulated with 1nM eODGT7 or eODGT8 np's for 24h. Black background depicts B-cells IL10Rα expression. Note the larger fraction of individual B-cells with expression of these cytokine pathway genes in eODGT8 np-treated cultured splenocytes relative to those treated with eODGT7 np's. **(B)** Table summary of all significantly upregulated genes in putative Breg/B10 B-cells (Loupe browser data combined across both eoDGT np treatment groups) relative to IL10ra non-expressing B-cells, ranked by log fold increase and p-values. Relevant B10-associated upregulated genes include CCL5 (RANTES), CD80, CxCl16 and Ccr2.

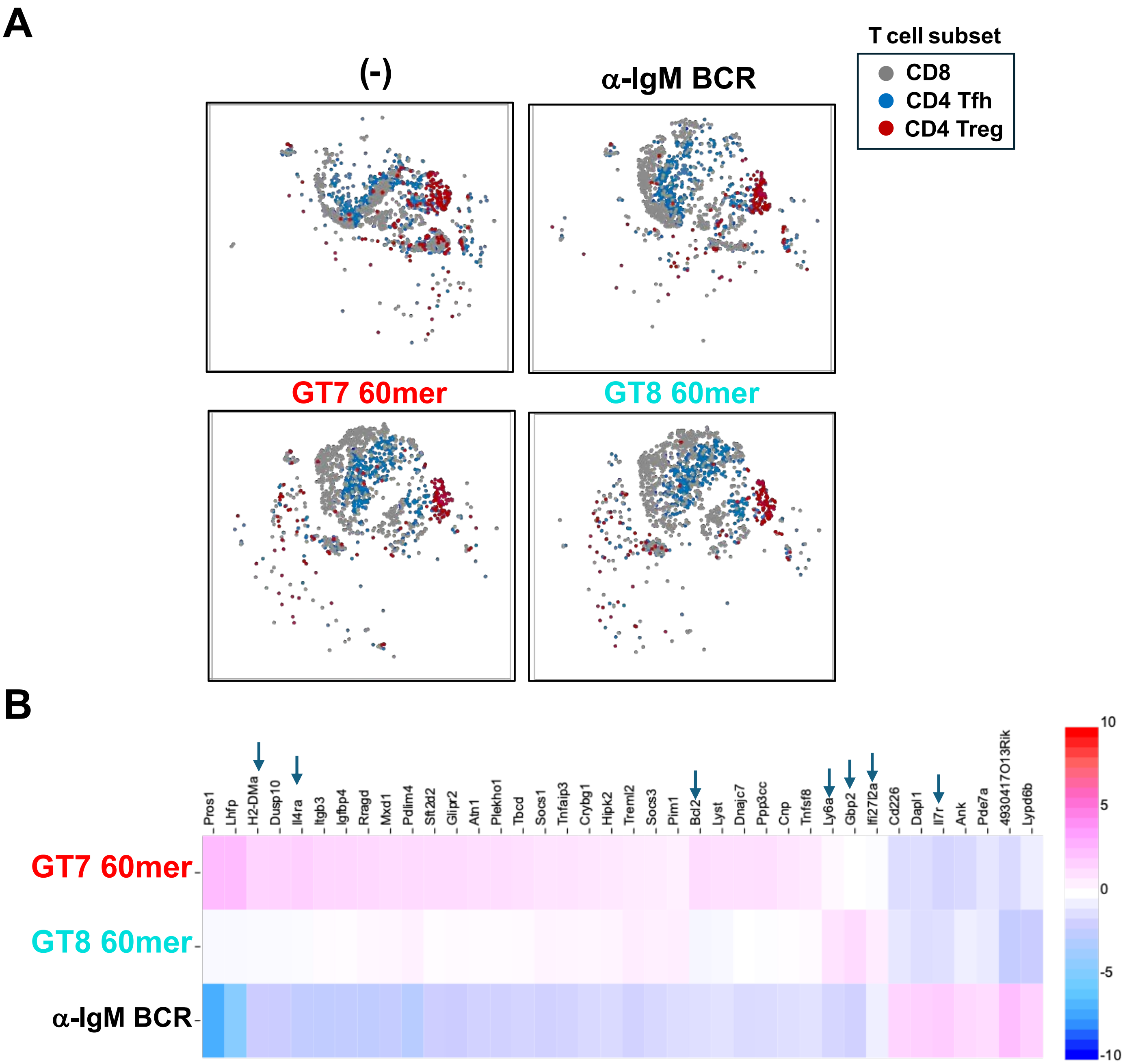


**Figure S9. eODGT7 and eODGT8 np-stimulated CH31 UCA$^{hom/hom\ dKI}$ splenic cultures exhibit expansion of CD4 T-cells with distinct Tfh transcriptional profiles.(A)** Gene expression maps showing distribution of T-cell subsets in all four treatment groups, with the $CD4^{+}$ Tfh-cell main zone (most in cluster 4) shown in blue, the CD8 T-cell main zone (most in cluster 3) shown in gray, and the CD4+ Treg-cell main zone (most in cluster 9) depicted in drak red. Multiple genes were found to be differentially expressed among CD4+T cells in the main zone (most in cluster 3; in blue) from the various treatments (see below panel B), while no significant differences were detected in the Treg population among treatments. **(B)** Heat maps of genes in main zone (cluster 3) CD4+ Tfh-cells exhibiting genes with significant differential expression (denoted by arrows, $p<0.05$ by Loupe Browser analysis) between eODGT7 and eODGT8 np-treated CH31UCAhom/homdKI cultures, relative to anti-BCR treatment. Analysis done this way shows that not only do eODGT8-treated splenic CD4+ T-cells tend to not only have decreased expression of Tfh2-like genes, have global decreased expression of multiple genes (or at least significant upregulation of genes in these cells was not observed compared to eODGT7), except for the three interferon-inducible genes associated with Tfh/$T_H2$ dampening.

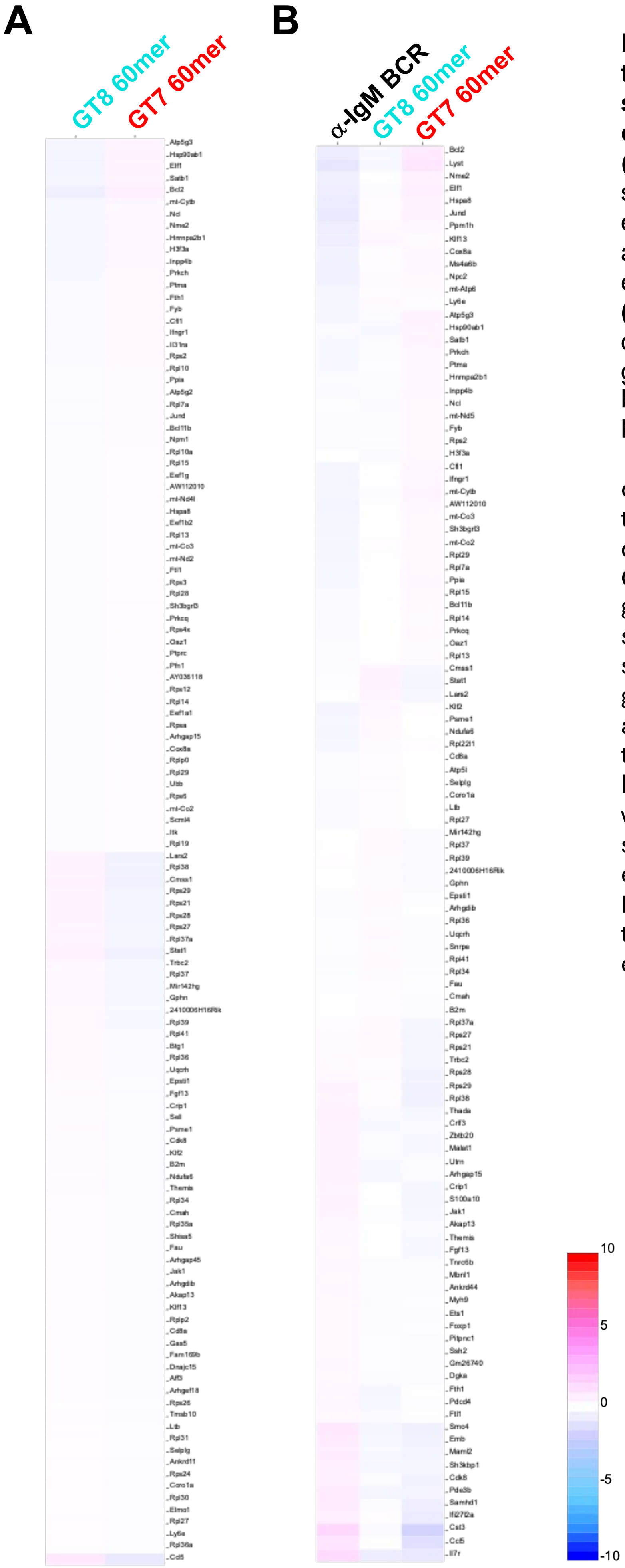


**Figure S10. Single CD8+ T-cell transcriptional profiles in eODGT np-stimulated CH31 UCA$^{hom/hom\ dKI}$ splenic cultures.** Shown are heat maps of cluster 3 (CD8+) CH31 UCAhom/homdKI splenocytes cultured for 24h with 1 nM eODGT7 vs. eODGT8 np's for 24h, either as head-to-head comparisons **(A)** or both eODGT np treatments, relative to anti-BCR **(B)**. Indicated at the bottom right is a differential gene expression index with fold gene up- or downregulation represented both numerically (-10 to +10) and by brightness of red or blue color, respectively.

Note that unlike head-to-head comparisons of eODGT7 vs eODGT8 np-treated CD4$^+$ T cells, no significantly differentially expressed genes were seen in CD8+ T cells from these two treatment groups. Furthermore, when genes with significant average counts from CD8+ splenocytes in both eODGT np-treated groups were compared to those from the anti-BCR treated group, only IL7R was found to be significantly higher in the latter. However, among all genes (ie, even those with low average counts), similar genes as seen in CD4$^+$ T cells were upregulated in eODGT7 np-treated cultures, such as IL4ra, Pros1, and H2-Dma compared to both treatments, while GBP1 was upregulated in eODGT8 CD8$^+$ T cells .

| Gene Name/ Lymphocyte population | Percentage of Splenic Cultures (by treatment group) | | | |
|---|---|---|---|---|
| | eODGT7 60mer | eODGT7 60mer | Anti-BCR | nil (-) |
| Ki67 in all B | 8.0 | 5.1 | 1.2 | 1.4 |
| IL10Rα in all B | 3.0 | 5.3 | 15.0 | 17.1 |
| H2-Eβ in all B | 20.3 | 10.1 | 1.7 | 3.5 |
| H2-Aα in all B | 32.2 | 18.0 | 8.0 | 11.1 |
| H2-Aβ in all B | 18.4 | 7.9 | 2.0 | 2.5 |
| IL4Rα in all $CD4^+$ T | 42.3 | 23.6 | 17.1 | 16.0 |
| IL7Rα in all $CD8^+$ T | 36.1 | 40.2 | 74.3 | 83.1 |

**Figure S11. CD4 Tfh-dependent GC program in eODGT7 np-treated CH31 $^{UCAhom/hom\ dKI}$ cultures is also associated with CD4+ and GC B-cell expansion.** Summary table showing the fraction of total B, $CD4^+$ T or $CD8^+$ T cells in all CH31 $UCA^{hom/homdKI}$ splenic cultures expressing genes listed, by treatment groups. Note that all genes listed had significant differences in gene expression ($p<0.05$ by Loupe Browser) between eODGT7 np & eODGT8 np treatments.  Also noteworthy is that the fraction of all listed genes in eODGT8 np-treated splenocytes yielded was an intermediate value between eODGT7 np treatment and the non-paratopic T-independent/innate control anti-BCR treatments.

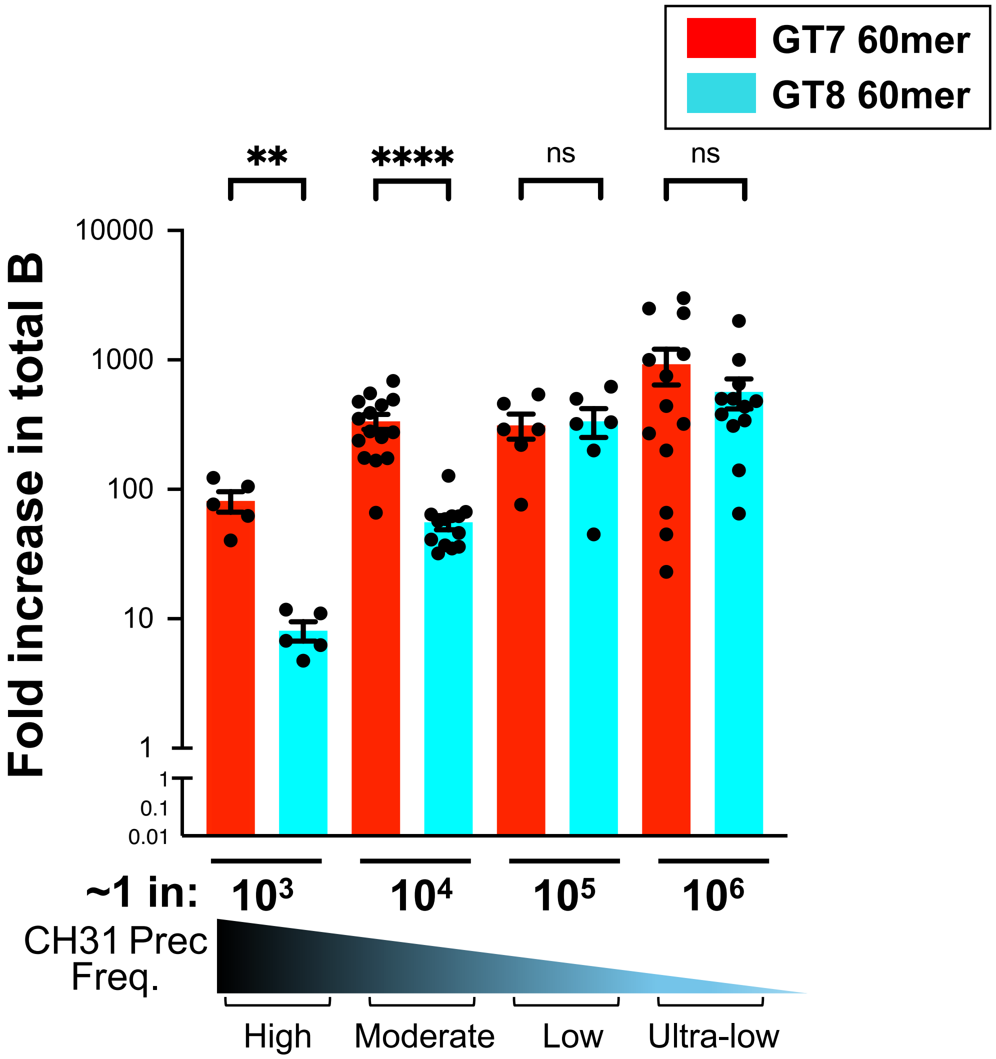


**Figure S12. Effect of precursor frequency on eODGT7 versus eODGT8 np abilities to clonally expand CH31 precursors.** Graphical representation of flow cytometrically-determined CH31$^{UCA\ hom/hom\ dKI}$ clonal expansion in spleens of CH31 UCA$^{hom/hom\ dKI}$→WT chimeras primed for 8d with eODGT7 or eODGT8 60mers. Clonal expansion was defined by flow gating as the fold increase in total donor-derived, on-target B-cells (live singlet, B220$^+$CD19$^+$ CD45.2$^+$ eODGT8 probe+ splenocytes) 8d post-prime, relative to original (pre-priming) reconstitution frequencies. Mann Whitney U tests were used to determine significance. *=p<0.05; **=p<.01, ***=p<0.001, ****=p<0.0001, ns=not significant.

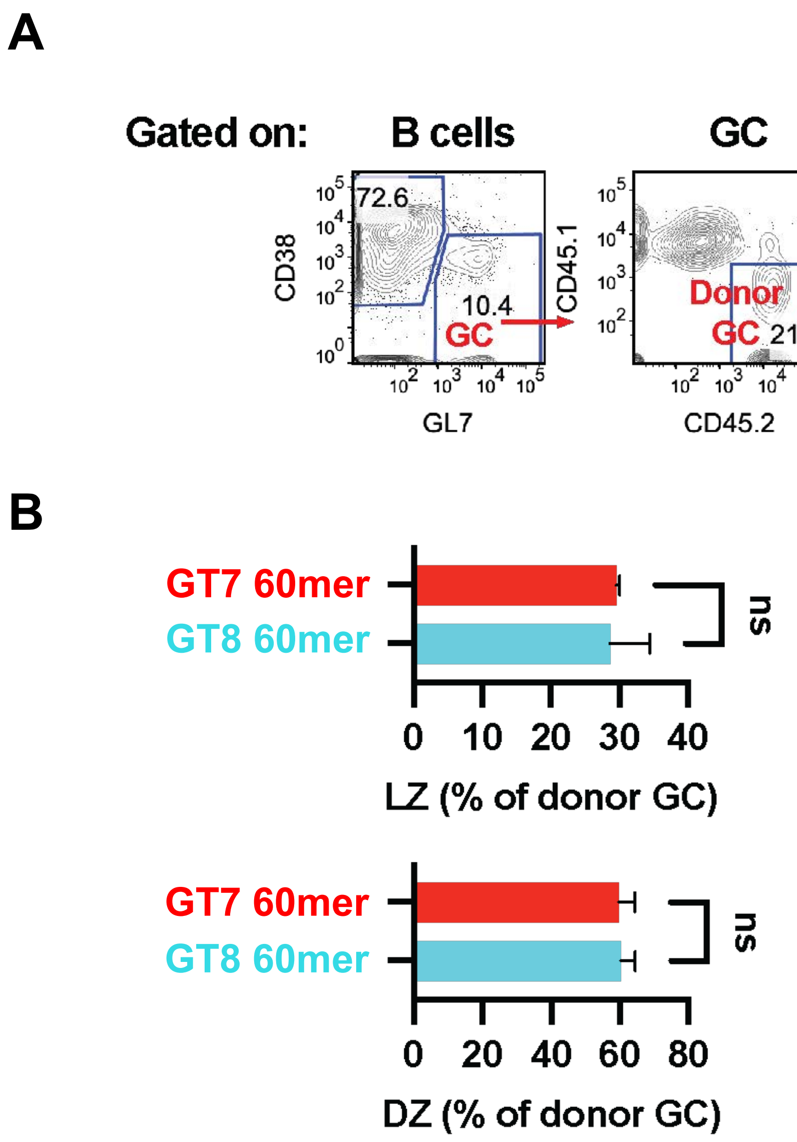


**Figure S13. Comparison of splenic CH31 UCA hom/hom dKI centroblast (CB) and centrocyte (CC) distributions in eODGT7 or eODGT8 np-primed CH31/WT chimeras.** Chimeras were reconstituted with low/physiologic (~1/10$^5$) frequencies of CH31 precursors and primed for 8 days with eODGT7 or eODGT8 60mers. **(A)** Representative FACS histogram from an eODGT-primed animal showing gating scheme to fractionate total (live, CD19,B220+) splenic B-cells into donor-specific (CD45.2+) centroblasts; CB (in GC Dark Zones; DZ) and centrocytes; CC (in GC Light Zones; LZ). **(B)** Graphical representation of CB and Cc percentages (+/-SD) found in eOD np-primed animals (*n*=5 per prime group). Mann Whitney U tests were used to determine significance. ns=not significant.

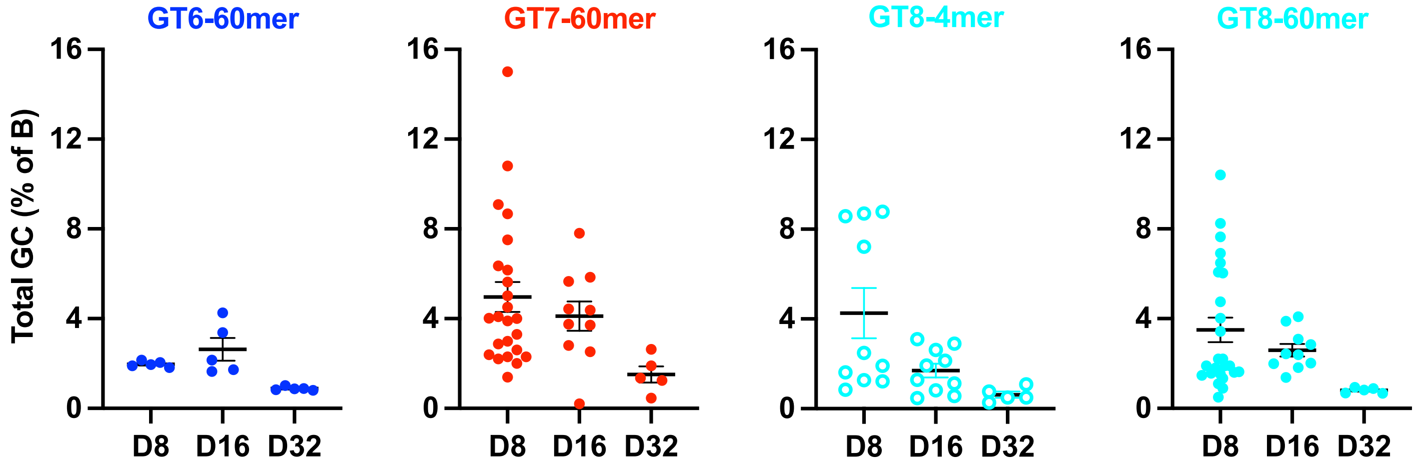


**Figure S14. Comparison of total GC formation in CH31/WT chimeras primed with eODGT immunogens for various durations.** Shown are graphical summaries of flow cytometric analysis of splenic GC+ B-cell frequencies in primed CH31 UCA$^{hom/hom\ dKI}$→WT chimeric animals, gated as the percentage of GC B-cells (CD38-GL7+) within live singlet, B220+CD19+ splenocytes (total B-cells). Chimeras were reconstituted with low/physiologic (~1/$10^5$) frequencies of CH31 precursors and primed for 8, 16, or 32 days with high valency eODGT6, eODGT7 or eODGT8 60mers, or (shown for reference), with low valency eODGT8 4mers. Each dot represents a primed chimeric animal, and black bars represent group means.

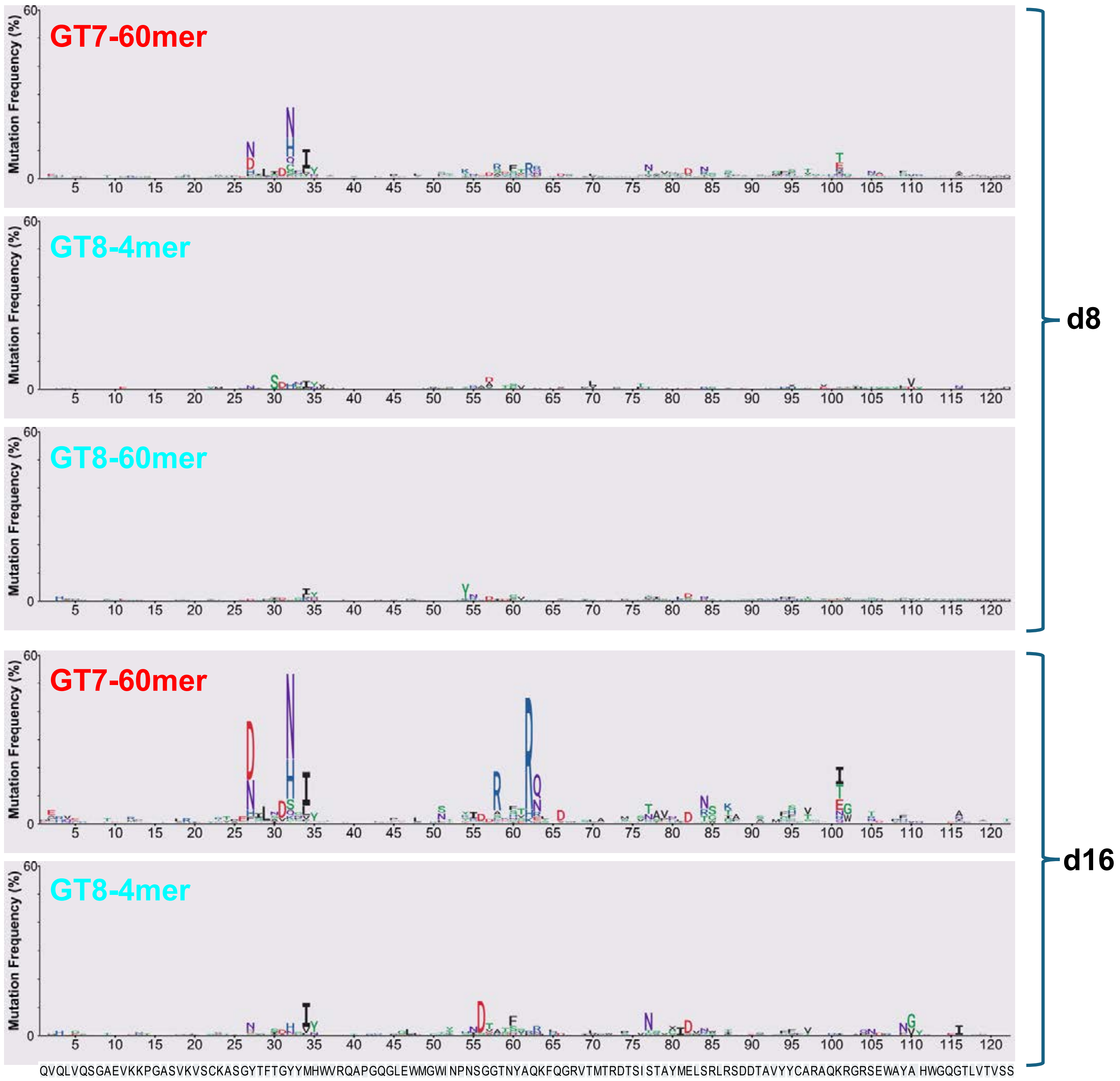


**Figure S15. Effect of 7th and 8th generation eODGT platform's differing $k_a/k_d$ tested at low/high valency on magnitude and duration of CH31 precursor V(D)J SHM induction.** Shown is the frequency of observed CH31 UCA HC rearrangement aa mutations, plotted as Logo plots of % residue changes by position shown on x-axis, amongst all bona fide CH31 UCA IgH/L pairs. CH31/WT Chimeras were reconstituted with low/physiologic (~$1/10^5$) frequencies of CH31 precursors and primed for 8 or 16 days with high valency eODGT7 or eODGT8 60mers, or low valency eODGT8 4mers. Individual total (unbiased) GC+ B-cells were recovered by the combination of flow cytometric sorting and 10X paired H/L seq, as described in the methods. A minimum of 1000 unique (non-oligoclonal), CH31 UCA V(D)J rearrangement HC/LC pairs from a minimum of five pooled animals per immunogen prime group were sampled by 10X-NGS. Note that data for the eODGT8 60mer-primed group is not shown at day 16 because absolute CH31 precursor numbers retained in GCs at this time point was too low for statistically meaningful comparison.

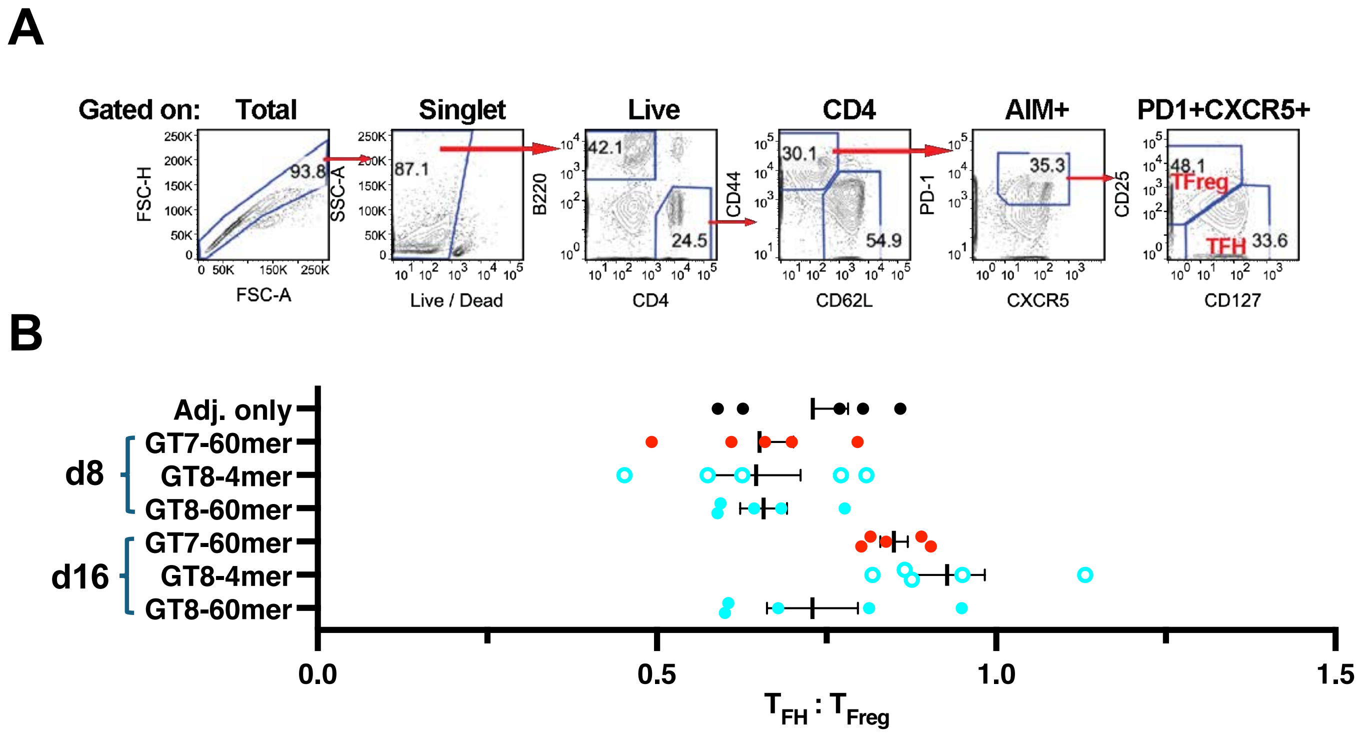


**Figure S16. Comparison of T follicular helper ($T_{FH}$) to T follicular regulatory ($TF_{Reg}$) CD4+ T cell ratios in eODGT-primed CH31/WT chimeras.** CH31 UCA$^{hom/hom\ dKI}$-->WT chimeras were generated by reconstitution with low/physiologic (~1/$10^5$) frequencies of CH31 precursors and primed for 8 or 16 days with either eODGT7 or eODGT8 high valency (60mer) np's, or low valency (tetrameric) eODGT8. **(A)** Representative FACS histogram from one eODGT-primed chimeric animal showing gating scheme to fractionate $T_{FH}$ or $TF_{Reg}$, gated as singlet, live, CD4+CD44+CD62L-PD1+CXCR5+ splenocytes that were CD25-CD127+ or CD25+CD127-, respectively. **(B)** Graphical summary of $T_{FH}$:$T_{Reg}$ ratios at peak splenic GC occupancy (d8) or peak SHM accumulation (d16) in CH31 UCA$^{hom/hom\ dKI}$→WT chimeric animals primed with eODGT7 np, eODGT8 np, and eODGT8 tetramers. Each dot represents a primed chimeric animal, and black bars represent group means (+/- SEM).
